# Validation of six commercial antibodies for detection of heterologous and endogenous TRPM8 ion channel expression

**DOI:** 10.1101/2022.11.29.518405

**Authors:** Pablo Hernández-Ortego, Remedios Torres-Montero, Elvira de la Peña, Félix Viana, Jorge Fernández-Trillo

## Abstract

TRPM8 is a non-selective cation channel expressed in primary sensory neurons and other tissues, including prostate and urothelium. Its participation in different physiological and pathological processes such as thermoregulation, pain, itch, inflammation and cancer has been widely described, making it a promising target for therapeutic approaches. The detection and quantification of TRPM8 seems crucial for advancing in the knowledge of the mechanisms under-lying its role in these pathophysiological conditions. Antibody-based techniques are commonly used for protein detection and quantification, although their performance with many ion channels, including TRPM8, is suboptimal. Thus, the search for reliable antibodies is of utmost importance. In this study, we characterized the performance of six TRPM8 commercial antibodies in three immunodetection techniques: western blot, immunocytochemistry and immunohistochemistry. Different outcomes were obtained for the tested antibodies; two of them proved to be successful detecting TRPM8 in the three approaches while, in the conditions tested, the other four were acceptable only for specific techniques. Considering our results, we offer some insight into the usefulness of these antibodies for detection of TRMP8 depending on the methodology of choice.

## Introduction

The detection of environmental temperatures is one of the critical functions of the somatosensory system. Transient Receptor Potential Melastatin 8 (TRMP8) is a non-selective cation channel that permeates divalent (Ca2+) and to a lesser extent monovalent (Na+, K+) cations [1,2]. It is a polymodal channel that can integrate different physical and chemical stimuli, being activated by mild cold and different natural and synthetic cooling compounds such as menthol, WS-12 [3] and icilin [4]. In addition, its activity is modulated by intracellular signaling molecules such as Ca2+ and PIP2 [5–7].

TRPM8 is preferentially expressed in Aɗ and C-fibers innervating the skin where it acts as a mild cold temperature transducer [8]. The depolarization of cold thermoreceptor endings generates trains of action potentials that travel from the periphery to-wards the cell body in the dorsal root (DRG) or trigeminal ganglia (TG), connecting with dorsal horn interneurons and to other brain regions (e.g. hypothalamus, cortex) where ultimately cold perception occurs and behavioral and autonomic thermoregulatory responses are triggered.

TRPM8 is also expressed in other surface tissues such as the cornea, where it participates in the regulation of the humidity of the eye surface by detection of the tear osmolality. Activation of corneal TRPM8+ endings triggers basal tearing and blinking, and its malfunction is implicated in the mechanisms of dry eye disease (DED) [9–13].

Expression of TRPM8 has also been described in tissues that are not exposed to the environment such as the brown adipose tissue where it has a role in thermogenesis and high-fat-diet induced obesity [14], intestinal epithelium associated with irritable bowel syndrome (IBS) and colitis [15– 17], and bladder associated with cooling-reflex, urinary urgency, overactive bladder and painful bladder syndrome [18–21]. Recently, TRPM8 expression has been detected in some brain regions (hypothalamus, septum, thalamus) and the retina, suggesting a role of this channel in thermal regulation and circadian control [22,23]. Additionally, TRPM8 is also expressed in prostate cancer cells. Some studies suggest a role of this channel in cell proliferation, whereas other findings suggest its participation in the reduction of metastatic processes in the prostate [24].

The important physiological role of TRPM8, as well as its involvement in different pathophysiological conditions (prostate cancer, migraine, obesity, cold pain, itch, inflammation, etc), makes this channel an important target for different studies. Currently, multiple techniques are used routinely to assess TRPM8 function. They include electrophysiological recordings and calcium imaging. Various mouse models are also available, including reporter and KO mice. In particular, reporter mice have been extremely useful in characterizing the expression pattern of TRPM8 in the mouse peripheral and central nervous system [22,25,26]. These studies, together with previous in-situ hybridization techniques [27], indicate that TRPM8 is expressed in a small sub-population of adult TG and DRG sensory neurons, and shows a restricted expression in the brain as well. In other species, the lack of reporter animals obliges the use of alter-native techniques, like immunofluorescence and western blot (WB) to quantify TRPM8 expression.

However, while immunofluorescence and WB are antibody-based mainstream techniques for protein visualization and quantification, the use of antibodies to characterize the expression of TRPM8, and other TRP channels, remains problematic [28]. There are many commercially available antibodies but their performance in different techniques like WB, immunocytochemistry (ICC) and immunohistochemistry (IHC) varies from acceptable to very poor, with many antibodies showing low specificity. To our knowledge, no systematic profiling of commercial antibodies against TRPM8 has been performed so far. Here, we validated six commercial TRPM8 antibodies for their use in WB, ICC and IHC, using different methodologies and following standard procedures. First, mouse TRPM8 fused to the fluorescence protein EYFP (mTRPM8-EYFP) was expressed in HEK-293 cells. ICC was performed in fixed cells testing the performance of the antibodies under TRPM8 overexpression conditions. In parallel, cell lysates were subjected to WB analysis with each of the tested antibodies using manufacturer recommended dilutions. Finally, the antibodies were tested against native TRPM8, using DRG sensory neurons from a mouse that specifically labels TRPM8+ neurons with EYFP. To confirm their specificity, the antibodies that performed best in DRG ICC and IHC were further validated using a TRPM8 KO mouse.

## Results

Six, commercially available, antibodies against TRPM8 were characterized in this study and summarized in Table 2 in the Material and Methods section. At present, five of them are still available and one has been discontinued (Origene1). All antibodies were designed against human TRPM8 but have reported cross-species reactivity with mouse and rat. As detailed in Table 2, two antibodies were polyclonal and four were monoclonal. The majority are directed against the extracellular domains of TRMP8 (although the exact region was only provided for Alomone) while Origene2 epitope is located in the N-terminal domain.

### All tested antibodies specifically label mouse TRPM8 overexpressed in HEK-293 cells

Cultured HEK-293 cells transfected with mTRPM8-EYFP were used to assess the suitability of each antibody in detecting overexpressed mTRPM8 by immunocytochemistry (ICC). EYFP fluorescence was used to identify transfected cells expressing TRPM8. First of all, using calcium imaging, we confirmed that EYFP+ HEK-293 cells responded to TRPM8 agonists such as mild cold stimuli (∼18-20 °C) and 10 μM WS-12, while EYFP-cells did not (Figure 1). Only cells that responded to 50 μM Carbachol, a compound that activates endogenous muscarinic receptors in HEK-293 cells [29], were included in the analysis.

**Figure 1.**
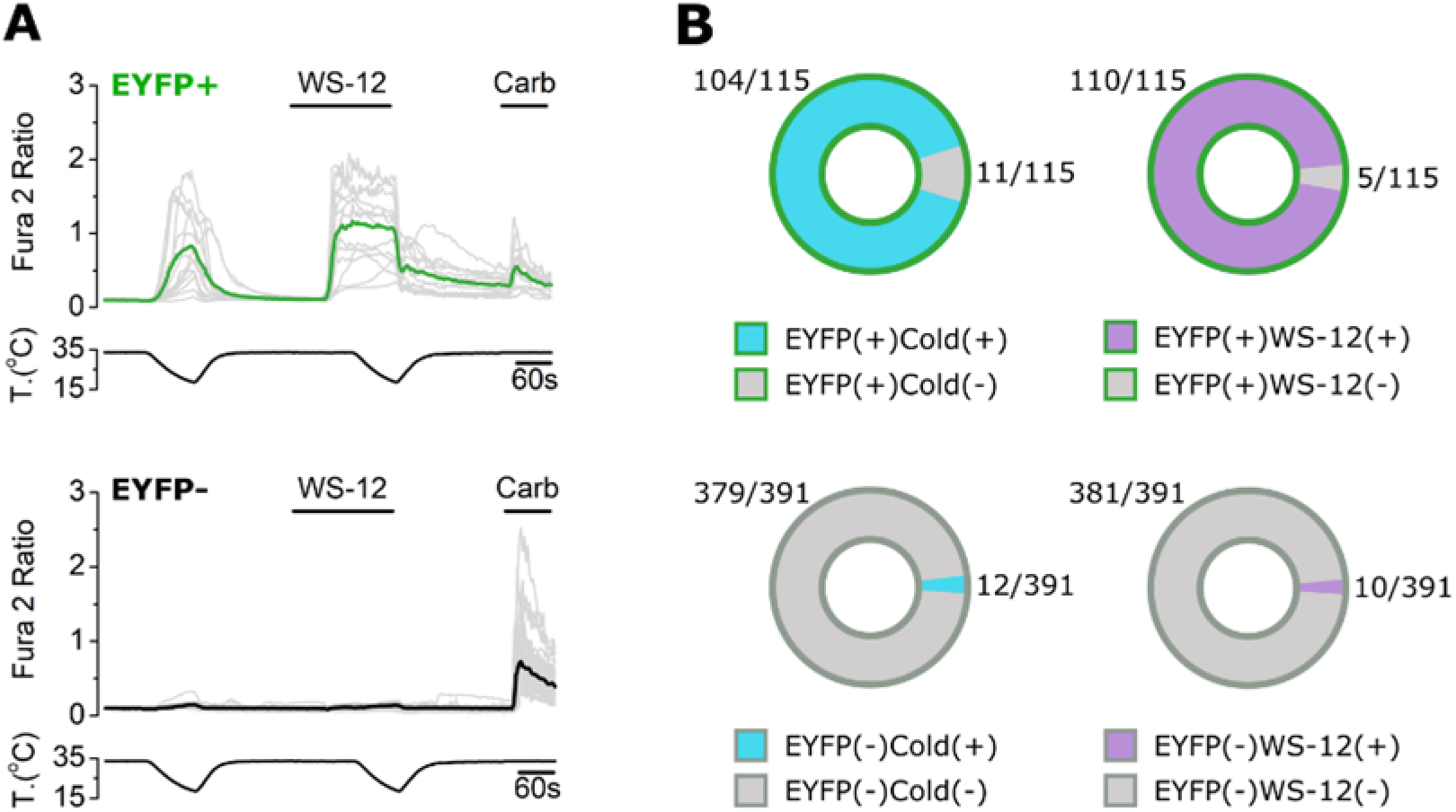
Calcium imaging in HEK-293 cells transiently overexpressing mTRPM8-EYFP. (A) Representative traces of calcium transients evoked by mild cold (∼18-20 °C), WS-12 (10 μM) and Carbachol (50 μM) in both EYFP+ and EYFP-cells. Traces corresponding to individual cells are shown in grey. Colored traces represent averages of the respective individual traces. (B) Proportion of EYFP+ and EYFP-cells responding to cold and WS-12. n = 506 cells; 2 independent experiments.

For mTRPM8 detection in ICC, we used two dilutions of each antibody in the range recommended by the vendors (1:500 or 1:200). All antibodies showed a clear co-localization of EYFP and mTRPM8 immunostaining (Figure 2A-F), indicating that the six antibodies are successfully detecting high levels of mTRPM8 protein. Notice the fluorescence punctate pattern typical of TRPM8 overexpression [30–32], both for the EYFP and antibody signal, further confirming antibody specificity (Figure S1). No TRPM8 immunostaining was found in neighboring EYFP- (i.e. non-transfected) cells, showing that none of the antibodies displayed any confounding unspecific staining. To compare the performance of the different antibodies, a specificity ratio (SR) (see Methods) was calculated for each dilution (Figure 2A-F, box plots). In general, the SR was similar with both dilutions or higher with 1:200 (ECM2 and Origene2), with the exception of ECM1 in which the higher dilution (1:500) showed a higher specificity ratio. In other words, ECM1 seems to work better when more diluted.

**Figure 2.**
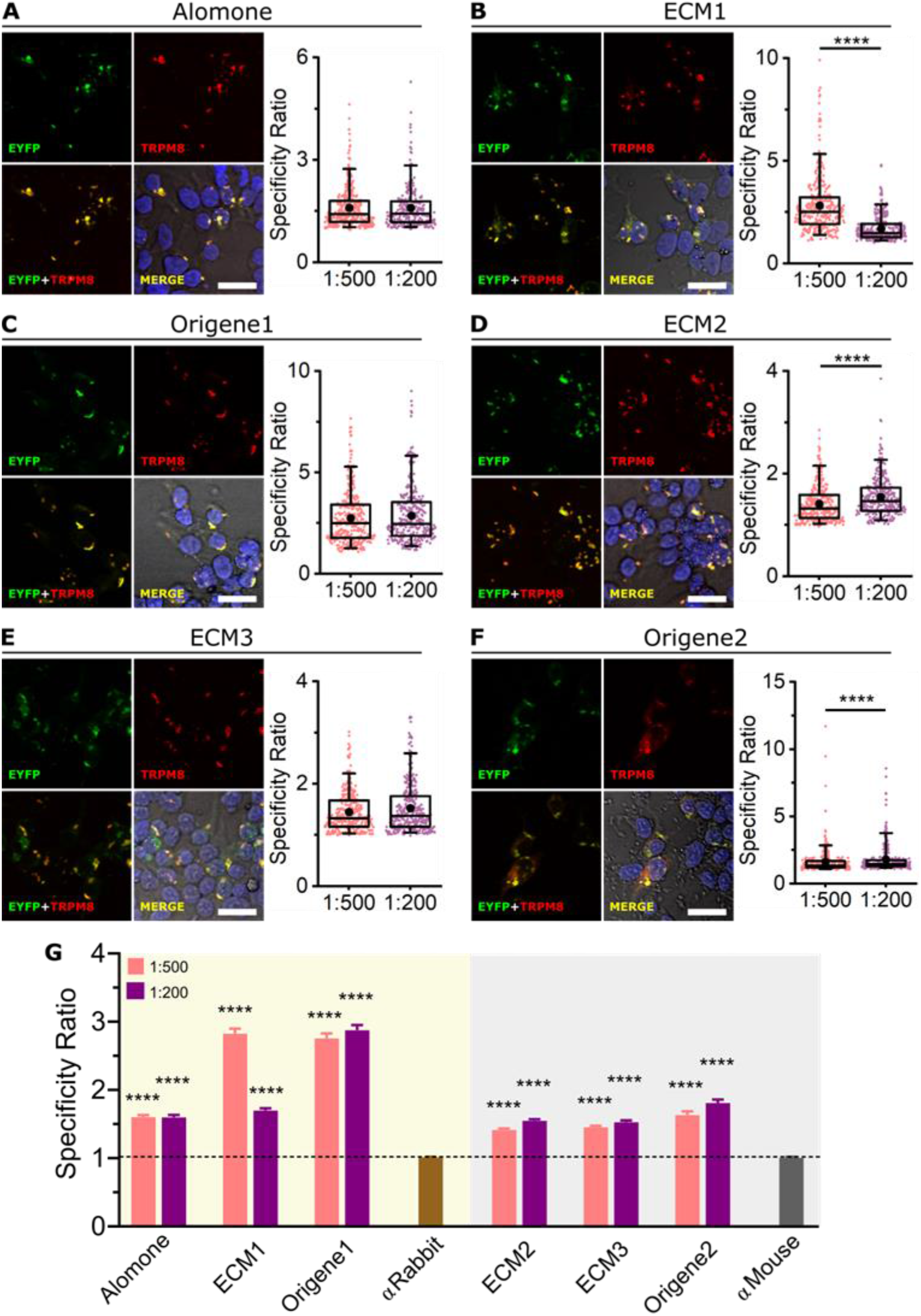
Immunocytochemistry of mTRPM8 in transfected HEK-293 cells. (A-F left) Confocal images of mTRPM8-EYFP transiently expressed in HEK-293 cell. EYFP (green), TRPM8 antibody (red) and Hoescht staining (blue). Merge image correspond to the overlap of the three fluorescent signals plus the bright field image. Scale bar: 30 μm. (A-F right) Box plots represent specificity ratio (SR) for each dilution of the respective antibody. Each dot corresponds to an individual EYFP+ cell. Box contains the 25th to 75th percentiles. Whiskers mark the 5th and 95th percentiles. The line inside the box denotes the median and the black dot represents the mean. (****p < 0.0001, Mann-Whitney test). (G) Bar histogram summarizing the SR mean ± SEM of each TRPM8 anti-body and the controls without primary antibody (αRabbit or αMouse secondary antibodies alone). Dashed line indicates SR = 1. (****p < 0.0001, Kruskal-Wallis followed by Dunn’s post hoc test vs αRabbit or αMouse). For each antibody and dilution, n > 270 cells; 4 fields from 2 independent transfections.

As shown in Figure 2G, the six antibodies have a SR significantly higher than the corresponding control without primary antibody (SR > 1, meaning that the antibody fluorescence signal is higher for TRPM8+ than for TRPM8-cells). However, SRs calculated using ECM1 and Origene1 were almost two-fold higher than with the other antibodies.

### Not all antibodies detect mTRPM8 in western blotting

Cell lysates from HEK-293 cells transfected with mTRPM8-EYFP were used to examine whether the mentioned antibodies were able to detect mTRPM8 in western blot (WB) (Figure 3A-F). A band with a molecular size around 160 kDa (black arrowheads), corresponding to mTRPM8-EYFP as expected, was detected by ECM1, Origene1 and Origene2. However, this band was not detected by Alomone, ECM2 or ECM3, even though EYFP immunoblotting revealed the presence of mTRPM8-EYFP in the sample (green arrowheads).

**Figure 3.**
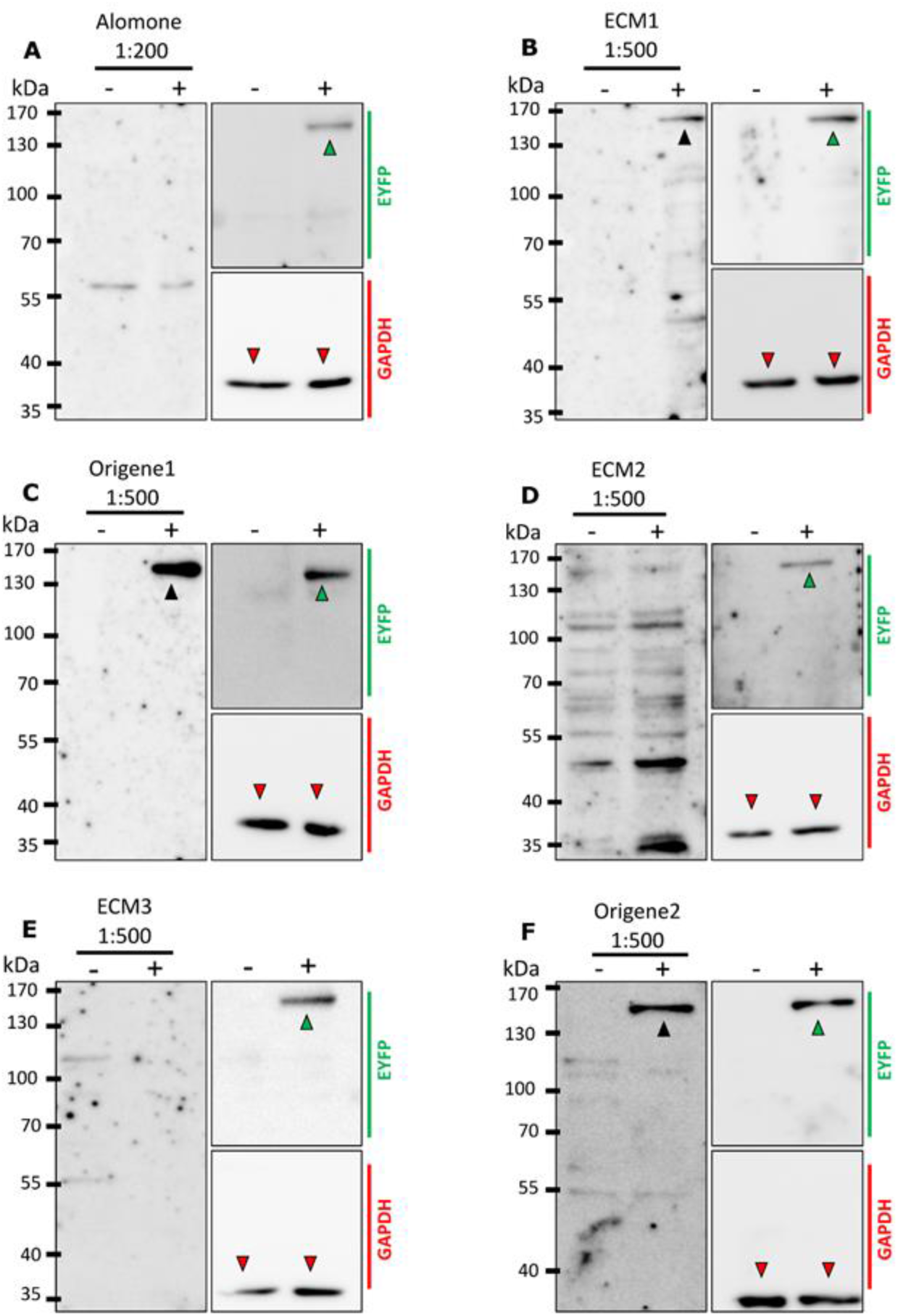
Western Blot analysis for TRPM8 antibodies specificity. (A-F) TRPM8 immunoblots using (A) Alomone (B) ECM1, (C) Origene1, (D) ECM2, (E) ECM3, or (F) Origene2 antibodies. (–) lanes: untransfected HEK-293 cells, (+) lanes: HEK-293 cells transfected with mTRPM8-EYFP. Left: Immunoblot with each TRPM8 antibody. Right-top: EYFP immunblotting on the same membrane. Right-bottom: GAPDH loading control. Black arrowheads indicate mTRPM8-EYFP bands revealed with antiTRPM8 antibody. Green arrowheads indicate mTRPM8-EYFP bands revealed with anti-GFP antibody. Red arrowheads indicate GAPDH bands. All blots were repeated at least 3 times to exclude a technical artefact when no anti-TRPM8 signal was observed. For each replicate, the same lysate was used for all antibodies.

### ECM1 and Origene1 specifically stain endogenous TRPM8 in mouse dorsal root ganglion sensory neurons

The specificity of the six antibodies against endogenously expressed TRPM8 was assessed by ICC and immunohistochemistry (IHC) in mouse dorsal root ganglion (DRG) neurons. These cells are first order sensory neurons that respond to diverse stimuli and include a subpopulation expressing TRPM8 that are activated by mild-cold and WS-12. In order to identify TRPM8+ neurons, we used a transgenic mouse line that expresses EYFP under the TRPM8 promoter (Trpm8BAC-EYFP+). Note that in this case EYFP is distributed freely in the cytoplasm and is not indicative of TRPM8 subcellular localization (Figure S1). Previous functional studies [33] indicated that, in this mouse line, EYFP expression is a good proxy of TRPM8 expression. Here, we confirmed the suitability of this reporter mouse to identify TRPM8-expressing neurons. Using cellular calcium imaging (Figure 4A, B), we found that the majority of cultured DRG cells with strong EYFP signal responded to mild cold (∼18-20 °C) and 10 μM WS-12 (100 % and 92.3 % respectively) while only few non- or dimly-fluorescent cells responded (2.37 % and 1.05 %). Only cells responding to high KCl (30 mM) were analyzed.

**Figure 4.**
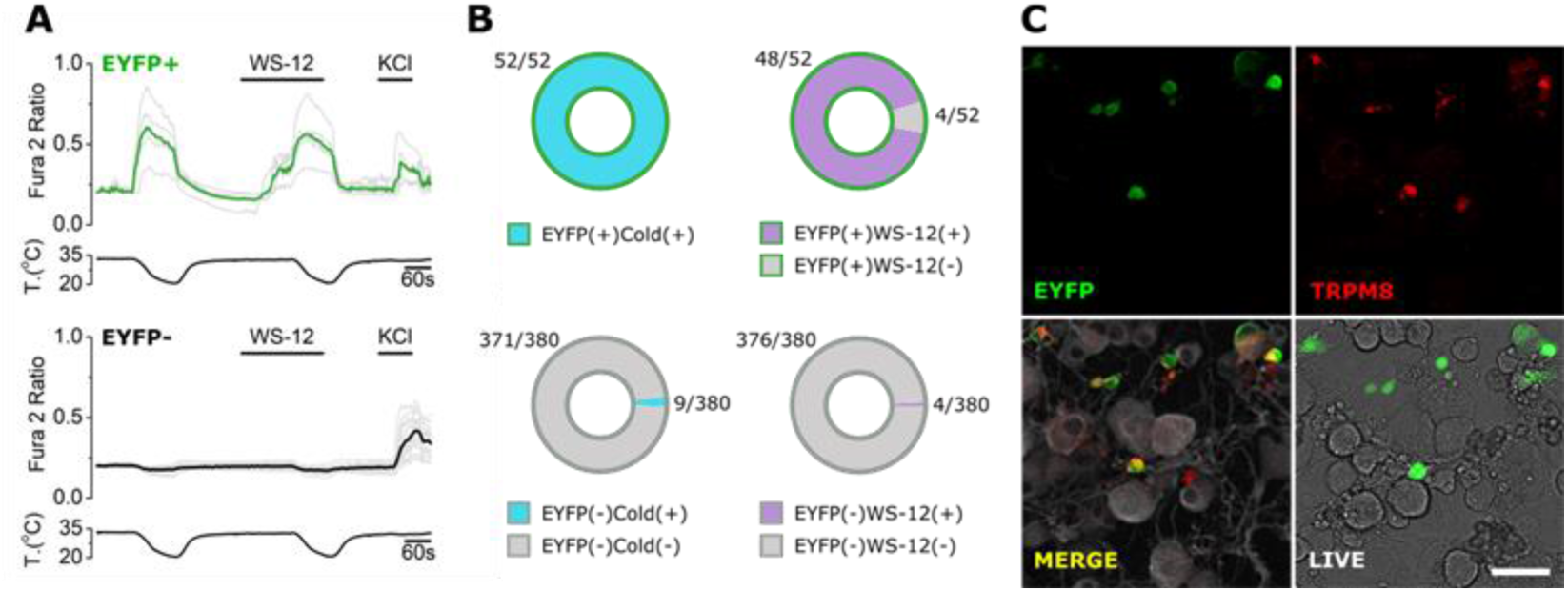
Calcium imaging in DRG cultured cells from Trpm8BAC-EYFP+ mice. (A) Representative traces of calcium transients evoked by mild cold (∼18-20 °C), WS-12 (10 μM) and KCl (30 mM) in EYFP+ and EYFP-neurons. Traces corresponding to individual cells are shown in grey. Colored traces represent averages of the respective individual traces. (B) Proportion of EYFP+ and EYFP-cells responding to cold and WS-12. (C) Representative immunocytochemistry performed after a calcium recording. Upper and bottom-left: confocal images of the immunofluorescence. Green: EYFP. Red: TRPM8 (ECM1 antibody). Grey: βIII-Tubulin. Bottom-right: transmitted light and EYFP fluorescence from the same live cells before fixation. n = 432 cells from 2 mice. Scale bar: 50 μm.

All six antibodies were tested for ICC on cultured DRG sensory neurons from Trpm8BAC-EYFP+ mice. Two recommended antibody dilutions (1:500 and 1:200) were used. Only two of the six antibodies performed well; ECM1 and Origene1 antibodies stained specifically the EYFP+ cold responsive cells (Figure 4C, 5B, C), showing no clear signal in EYFP-neurons, and displayed a SR > 1 (Figure 5B, C, G). In our conditions, the other four antibodies performed poorly. Alomone stained almost all sensory neurons (whether EYFP+ or EYFP-, Figure 5A), which was inconsistent with the restricted expression of TRPM8. In contrast, ECM2 did not label any cell (Figure 5D). ECM3 stained some cells but it was not specific for EYFP+ neurons (Figure 5E). Origene2 seemed to mark EYFP+ cells with more intensity than EYFP-but showed some unspecific staining (Figure 5F).

**Figure 5.**
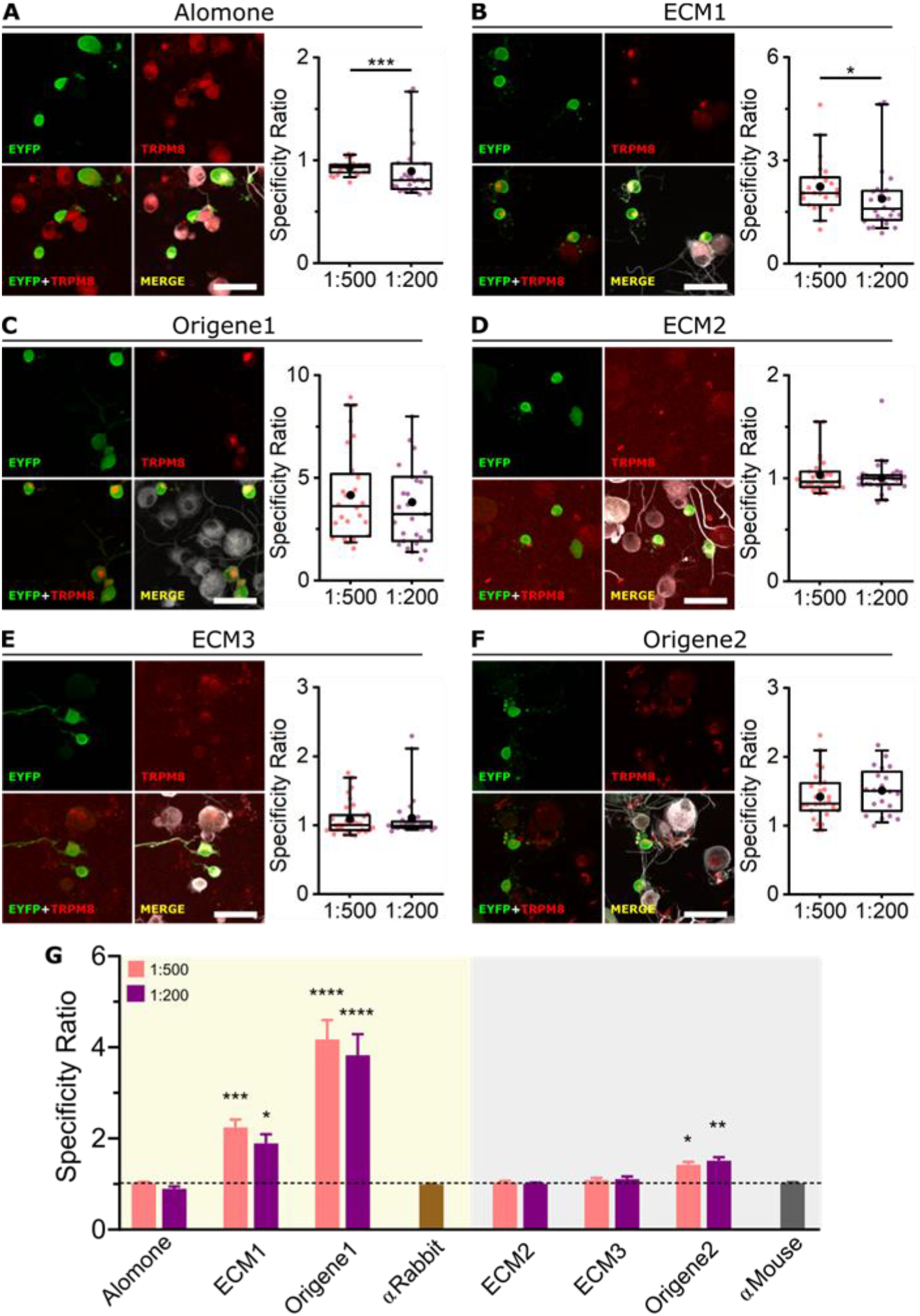
Immunocytochemistry of endogenous TRPM8 in cultured DRG cells from the Trpm8BAC-EYFP+ mouse. (A-F left) Confocal images of TRPM8+ sensory neurons. EYFP (green), TRPM8 antibody (red) and βIII-Tubulin (grey). Scale bar: 50 μm. (A-F right) Box plots represent specificity ratio (SR) for each dilution of the respective antibody. Each dot corresponds to an individual EYFP+ cell. Box contains the 25th to 75th percentiles. Whiskers mark the 5th and 95th percentiles. The line inside the box denotes the median and the black dot represents the mean. (*p < 0.05, ***p < 0.001, Mann-Whitney test). (G) Bar histogram summarizing the SR mean ± SEM of each TRPM8 antibody and the controls without primary antibody (αRabbit or αMouse secondary antibodies alone). Dashed line indicates SR = 1. (*p < 0.05, **p < 0.01, ***p < 0.001, ****p < 0.0001, Kruskal-Wallis followed by Dunn’s post hoc test vs αRabbit or αMouse). n = at least 20 cells; 4 pictures from 2 mice for each antibody and dilution.

The antibody concentrations used did not seem to have an important effect in their SR, with the exception of Alomone and ECM1, which showed a higher SR with 1:500 dilution. (Figure 5A-F, box plots). Comparing the SR of each antibody versus the control (no primary antibody) (Figure 5G) we observed that ECM1 and Origene1 have a substantially higher ratio suggesting a good specificity. Origene2 SR is slightly higher than the control indicating that this antibody also shows some specificity for TRMP8 in the conditions used.

Endogenous protein expression is more commonly assessed with IHC than with ICC, as the sample is more easily acquired and tissue structure is preserved. However, other challenges arise such as epitope accessibility and masking [34,35]. We tested all six antibodies in IHC of mouse DRG slices. Initially, citrate antigen retrieval was used, but neither antibody worked properly; subsequent IHCs were performed in the absence of antigen retrieval. The same two dilutions (1:500 and 1:200) were used, without any significant difference in the SR between them (Figure 6A-F, box plots). Similar to the ICCs, only ECM1 and Origene1 showed staining restricted to the bright EYFP+ neurons and distinguishable from the unspecific signal (Figure 6B, C). Alike the ICC results, Alomone marked all cells unspecifically (Figure 6A), and ECM2 and ECM3 produced an unspecific staining pattern similar to the staining observed with the secondary antibody alone (Figs. 6D, E, S2). In addition, some EYFP-cells were strongly labelled after ECM2 incubation. With Origene2, EYFP+ cells were immunostained but a fraction of the EYFP-cells was also unspecifically labeled.

**Figure 6.**
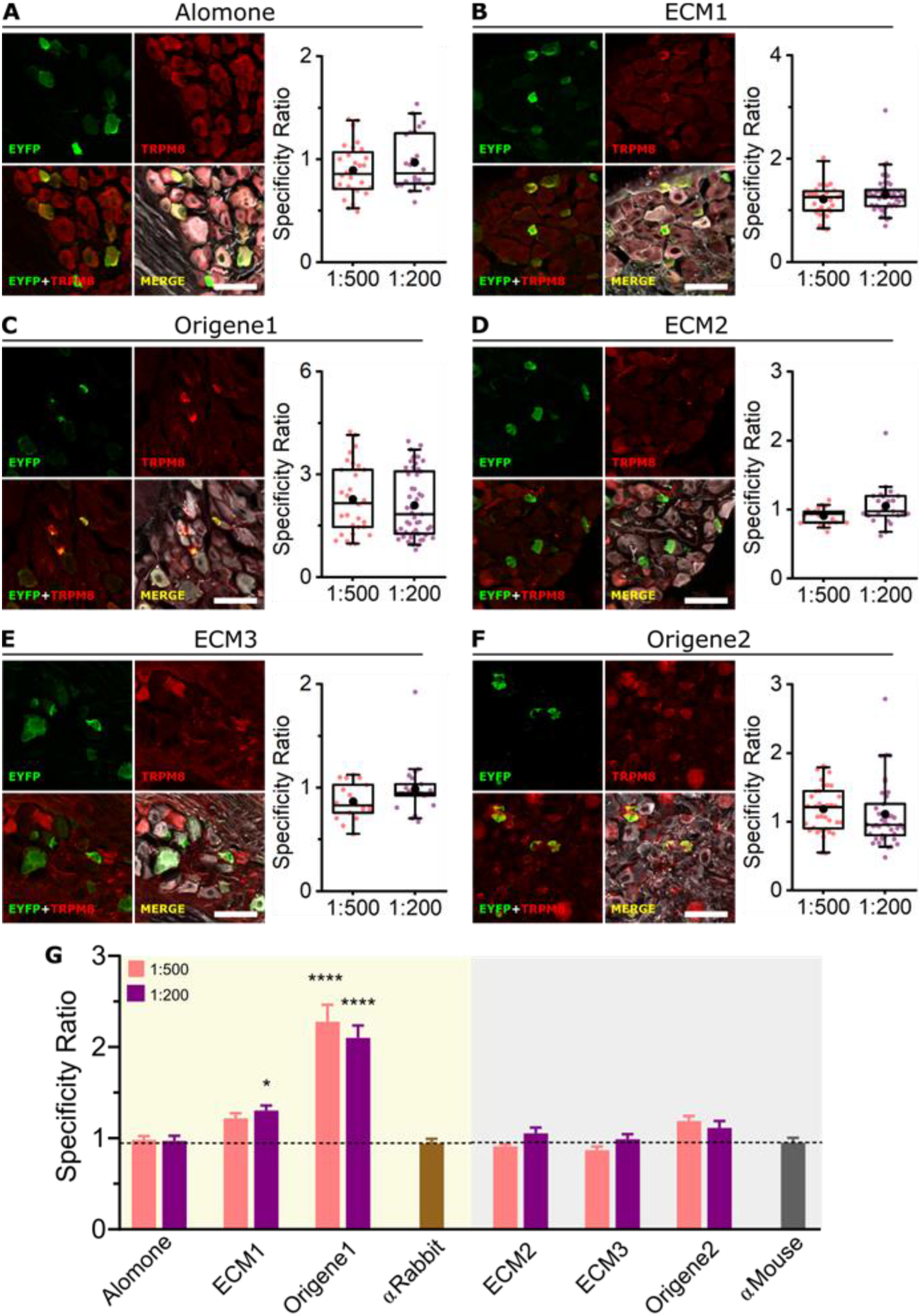
Immunohistochemistry of endogenous TRPM8 in DRG slices from the Trpm8BAC-EYFP+ mouse. (A-F left) Confocal images of TRPM8+ sensory neurons. EYFP (green), TRPM8 antibody (red) and βIII-Tubulin (grey). Scale bar: 50 μm. (A-F right) Box plots represent specificity ratio (SR) for each dilution of the respective antibody. Each dot corresponds to an individual EYFP+ cell. Box contains the 25th to 75th percentiles. Whiskers mark the 5th and 95th percentiles. The line inside the box denotes the median and the black dot represents the mean. (Differences among dilutions were not significant, Mann-Whitney test). (G) Bar histogram summarizing the SR mean ± SEM of each TRPM8 antibody and the controls without primary antibody (αRabbit or αMouse secondary antibodies alone). Dashed line indicates SR = 1. (*p < 0.05, ****p < 0.0001, Kruskal-Wallis followed by Dunn’s post hoc test vs αRabbit or αMouse). n = at least 18 cells; 4 images from 2 mice for each antibody and dilution.

The SR comparison among the different antibodies is shown in Figure 6G: only ECM1 (1:200 dilution) and Origene1 have a SR different from the control without primary antibody.

### Validation of TRPM8 antibody specificity in KO animals

Knockout cells provide the best negative control in antibody validation [36]. Both, the ICC and IHC results suggest that only ECM1 and Origene1 antibodies perform well in detecting endogenous expression of mTRPM8. To validate the specificity of these two antibodies, both ICC and IHC were repeated using DRG cultured cells and slices from TRPM8 KO mice (Trpm8EGFPf ; B6;129S1(FVB)-Trpm8tm1Apat/J) [25] that express farnesylated EGFP instead of TRPM8 (i.e. EGFP marks neurons that should have ex-pressed TRPM8). Using either ECM1 or Origene1, specific antibody fluorescence was absent in the EGFP+ cells of the TRPM8 KO mouse (Figures 7, S3). An unspecific signal was detected in some EGFP-cells (Figure 7A, B), albeit with a different fluorescence distribution and dimmer intensity than in EYFP+ cells in the TRPM8 reporter mouse (Trpm8BAC-EYFP+) (Figure S3). Furthermore, the SR for both antibodies was significantly higher in the reporter mouse compared to the TRPM8 KO both in ICC and IHC (Figure 7C, D).

**Figure 7.**
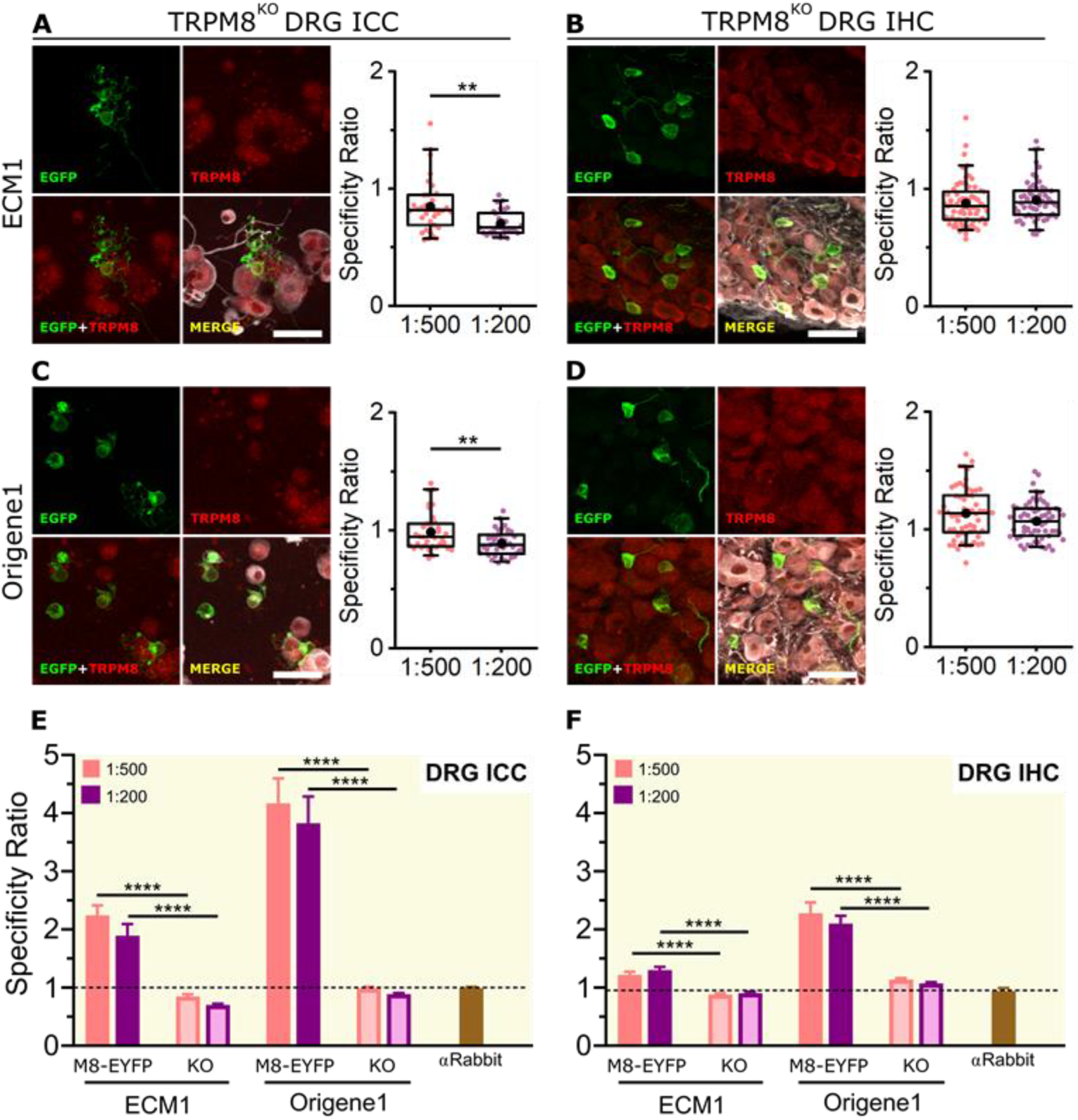
Immunofluorescence of endogenous TRPM8 in DRG cells and slices from the TRPM8 KO mouse. (A, C) Immunocytochemistry. (B, D) Immunohistochemistry. (A-D left) Confocal images of TRPM8 KO (Trpm8EGFPf; B6;129S1(FVB)-Trpm8tm1Apat/J) sensory neurons. EGFP (green), TRPM8 antibody (red), βIII-Tubulin (grey). Scale bar: 50 μm. (A-D right) Box plots represent specificity ratio (SR) for each dilution of the respective antibody. Each dot corresponds to an individual EGFP+ cell. Box contains the 25th to 75th percentiles. Whiskers mark the 5th and 95th percentiles. The line inside the box denotes the median and the black dot represents the mean. (**p < 0.01, Mann-Whitney test). (E, F) Bar histograms summarizing the SR mean ± SEM of each TRPM8 antibody in the KO and the reporter (Trpm8BAC-EYFP+, named M8-EYFP in the figure) mouse in ICC (E) and IHC (F). Dash line indicates mean SR for the control without primary antibody (****p < 0.0001, Mann-Whitney test). n = at least 22 cells; 4 pictures from 2 mice for each antibody and dilution.

## Discussion

The use of antibodies to quantify the level or expression pattern of antigens (e.g. proteins) is one of the most popular and valuable techniques in cell biology. It is com-mon knowledge that performance of many antibodies is substandard, with low sensitivity and poor specificity [36]. Unfortunately, very often, negative results are not reported in the literature. In other cases, antibody specificity is not assessed critically, which could lead to erroneous interpretation of the results. Thus, proper characterization of antibodies is of utmost importance [36,37].

Our main aim was to compare the performance of six commercial TRPM8 anti-bodies in western blot (WB), immunocytochemistry (ICC) and immunohistochemistry (IHC). Our results provide useful hints for choosing the most adequate antibodies for different TRPM8 protein detection techniques. In our hands, the six antibodies analyzed show diverse suitability for each technique and expression system. According to the respective manufacturers, all six antibodies should work for WB. However, using their recommended dilutions, only three of them yielded satisfactory results for our standards. Interestingly, Alomone antibody has been widely used for WB [38–40] but, in our conditions, it did not work properly. However, in many of these articles, the antibody dilution was lower than recommended and the reported TRPM8 bands were dubious and/or no full membrane/molecular weight markers were shown, making it difficult to rule out the presence of unspecific bands or cross-detection. In some cases, the reference to the antibody used was not clear; we could not rule out that it was a different antibody from the one we tested. Also, some studies targeted human TRPM8 [41,42] and we used the mouse ortholog instead. Alomone antibody immunogen is publicly known (Table 2); it comprises 13 residues of human TRPM8 from which 2 are different in human and mouse. This may explain the discrepancy between the results of this work and what others reported with human TRPM8. ECM2 and ECM3 also failed to produce a good WB signal. In one of the replicates with ECM2, we detected a faint band of a molecular weight compatible with TRPM8, but it is also present in the lysate from non-transfected cells, thus questioning its specificity. Many other unspecific bands of different molecular weights were also detected, so special caution has to be taken if using this antibody for WB (Figure S4). At the time of preparing this manuscript, to our knowledge, no published work references either ECM2 or ECM3 for WB. Interestingly, both ECM2 and ECM3, as well as Alomone, worked well in ICC of mTRPM8 transfected HEK-293 cells, suggesting that they are in fact able to bind to mTRPM8. The discrepancy observed between WB and ICC could be explained by the fact that cell lysis and denaturing conditions used in WB may make the epitope less accessible to the antibody paratope. For those reasons, a potential use of these three antibodies for WB after improving conditions cannot be excluded.

ECM1, Origene1 and Origene2 produced consistently good results for WB and ICC in a TRPM8 overexpression system, indicating that, at least in the conditions of this study, they can be confidently used for both techniques. Limited literature references Origene1 [43]. No previous WB records have been found for ECM1 or Origene2, making this study the first to show their appropriateness for WB.

Proteins are differentially expressed in heterologous and endogenous systems, being generally more abundant in the former. Endogenous TRPM8 detection was more challenging and only ECM1 and Origene1 worked well both in dorsal root ganglion (DRG) ICC and IHC. We cannot give a firm judgement about Origene2 as the immunostaining in DRG ICC was ambiguous. An additional consideration with Origene2 is that it was raised in mouse, thus the anti-mouse secondary antibody could bind to immunoglobulins present in mouse DRG distorting the interpretation of the results (Figure S2). Testing this antibody in human samples could help in resolving this issue. These results, together with its good performance in WB and ICC of HEK-293 cells, suggest that Origene2 could potentially be a good antibody for endogenous TRPM8 detection after adjusting experimental protocols.

In our hands, Alomone, ECM2 and ECM3 did not have enough sensitivity and/or specificity to detect endogenous TRPM8, at least with the conventional conditions we used. There is abundant literature using Alomone antibody for detection of endogenous TRPM8 by ICC and/or IHC in different tissues [39,44–48]. However, in many of the published figures, the staining pattern was not consistent with the known restrict-ed expression of TRPM8 [31,49]. In many cases, all cells seem to be stained with fluorescence evenly spread across the entire cell. In contrast, the fluorescence we see with antibodies that we consider good (ECM1 and Origene1) is distributed in a punctate pattern (Figure S1). Moreover, in some studies this antibody marks almost all cells subjected to immunochemistry. This is inconsistent with the known expression pattern of TRPM8 in DRG neurons, which is restricted to a small subpopulation of about 10-15 % cells. In the light of our results, we believe that caution should be taken interpreting the immunofluorescence signal from those studies. Similar to the reasons proposed for WB, we cannot exclude that this antibody could perform better in human tissue or in different experimental conditions.

Our data leads us to suggest the use of ECM1, Origene1 and Origene2 for WB and ECM1 and Origene1 for immunochemistry of endogenous TRPM8. A brief comparison of the performance of the tested antibodies with the different methodologies is shown in Table 1.

**Table 1.** Antibody performance with the different techniques used in this study. – poor, + regular, ++ good, +++ excellent.

| Antibody | Host | HEK-293 + TRPM8-EYFP |  | Mouse DRG |  |
| --- | --- | --- | --- | --- | --- |
|  |  | WB | ICC | ICC | IHC |
| Alomone | Rabbit | - | ++ | - | - |
| ECM1 | Rabbit | +++ | +++ | ++ | ++ |
| Origene1 | Rabbit | +++ | +++ | +++ | +++ |
| ECM2 | Mouse | - | ++ | - | - |
| ECM3 | Mouse | - | ++ | - | - |
| Origene2 | Mouse | +++ | ++ | + | - |

Origene1 has been discontinued. According to the available information, it seems to come from the same clone(EPR4196(2)) as Abcam ab109308 (also discontinued) that has been used with good results for WB and IHC [49–51]. We think that Abcam anti-body could still be available in many labs and could be a good alternative for TRPM8 detection.

### Limitations of the study

Variations in immunodetection protocols result in countless procedures in the literature. Variables include fixation, sample preparation and tissue processing, antibody concentration, antigen retrieval, permeabilization, blocking of non-specific sites, signal amplification, detection method and other [52]. There is no generic protocol for immunolabeling. We standardized the labeling procedure for the six antibodies tested but did not performed an in depth exploration of the protocol space. It is perfectly feasible that other conditions would have resulted in better results with some of the antibodies tested. Various online resources provide useful tips on step-by-step guides for staining optimization.

## Material and Methods

### Animals

All experimental procedures were performed in accordance with the Spanish Royal Decree 53/2013 and the European Community Council Directive 2010/63/EU. Adult mice (2-4 months old) of either sex were used. Mice were housed in a temperature-controlled room (21 °C) on a 12 h light/dark cycle, with access to food and water ad libitum.

### Mouse lines

Trpm8BAC-EYFP+ BAC transgenic line was generated in our laboratory: enhanced yellow fluorescent protein (EYFP) is expressed under the promoter of TRPM8 [33].

Trpm8EGFPf; B6;129S1(FVB)-Trpm8tm1Apat/J was obtained from Ardem Patapoutian (Scripps Research Institute) and expresses enhanced green fluorescent protein (EGFP) after the TRPM8 start codon; homozygous mice are null for TRPM8 [25]. The lox-P flanqued neomycin cassette introduced in the Trpm8 locus during the generation of the transgene was removed to enhance GFP expression [53]. The genotype of all mice was confirmed by PCR.

### Cell line culture and transfection

The HEK-293 cell line was obtained from ECACC (Salisbury, UK). Cells were maintained at 37 °C, 5% CO2 in DMEM medium (Invitrogen, USA) supplemented with 10% fetal bovine serum and 1% penicillin/streptomycin. 24h before transfection, cells were seeded on 6-well plates. For ICC, wells were filled with poly-L-lysine (0.01%, Sigma, USA) treated 6 mm diameter #0 glass coverslips. Cells were transfected with mTRPM8-EYFP (mTRPM8 fused with EYFP at its C-terminus) [54] using Lipofectami-neTM 2000 (Thermofisher Scientific, USA), 3 μg DNA and 3 μL Lipofectamine in 300 μl Optimem (Thermofisher Scientific, USA) were used per well containing 700 μl DMEM. 24-48h post-transfection, mTRPM8-EYFP expression was monitored with an epifluorescence microscope and cells were either lysated (for WB) or fixed in 4% PFA (for ICC).

### Mouse DRG extraction and neuronal culture

Mouse DRG neurons were extracted and dissociated as previously described [55]. Briefly, mice were euthanized by cervical dislocation. The spinal cord was removed and 20-40 DRGs were dissected and washed in cold HBSS solution (Invitrogen, USA). For IHC, whole ganglia were directly fixed for 2 h in 4 % PFA. For ICC, ganglia were then incubated in 900 U/mL type XI collagenase (Sigma, USA) and 5.46 U/mL dispase (Invitrogen, USA) for 45 min at 37 °C in 5% CO2. After enzymatic treatment, ganglia were mechanically dissociated using fire-polished glass pipettes in calcium-free solution containing HBSS (Invitrogen, USA), 1% MEM-Vit (Invitrogen, USA), 10% fetal bovine serum (Invitrogen, USA) and 100 mg/mL penicillin/streptomycin. The cell suspension was centrifuged and the pellet resuspended in culture medium containing: MEM (Invitrogen, USA), 1% MEM-Vit (Invitrogen, USA), 10% fetal bovine serum (Invitrogen, USA) and 100 mg/mL penicillin/streptomycin. Cells were then seeded on 6 mm diameter glass coverslips previously coated with 0.01% poly-L-lysine (Sigma, USA). 24h after seeding, cells were subjected to calcium imaging or fixed for 10 min in 4%PFA.

### Calcium microfluorometry

Fura2-AM calcium indicator (Thermo Fisher Scientific, USA) was used for ratiometric calcium imaging experiments. Cells were loaded with 5 μM Fura2-AM and 400 ng/mL Pluronic F-127 (Thermo Fisher Scientific, USA) in control external solution for 45 min at 37 °C in 5% CO2. Coverslips containing the cells were placed in a low volume chamber mounted on an inverted microscope (Leica DMI3000B, Leica Microsystems, Germany) and continuously perfused with fresh solution at a rate of ∼1ml/min. Fura2-AM was excited at 340 and 380 nm with a Lambda 10-2 filter wheel and a Lambda LS xenon arc lamp (Sutter Instruments, USA). Emission fluorescence was filtered with a 510 nm long-pass filter. Images were acquired with an Orca ER CCD camera (Hamamatsu Photonics, Japan) at a frequency of 0.33 Hz and analyzed with MetaFluor software (Molecular Devices, USA). Cytosolic calcium increases are presented as the ratio of emission intensities after sequential excitation at 340 and 380 nm (F340/F380). Measurements in HEK293-cells and DRG neurons were performed at 32-34 °C using a homemade water-cooled Peltier system controlled by a temperature feedback device.

### Western Blot

All WB experiments were done on samples obtained from HEK-293 cells transfected with mTRPM8-EYFP. Cells were harvested 48 h after transfection, centrifuged at 800G 10 min and washed with cold PBS twice. Pellets were lysed (Buffer Lysis: 50 mM Tris-HCl, pH 7.5, 120 mM NaCl, 0.5 mM EDTA, 0.5 % Nonidet P-40 with phosphatase, and protease inhibitors (cOmplete Mini, Roche)). Cell lysates were sonicated for 10 min at 4 °C, and centrifuged at 10,000G for 15 min at 4 °C. Protein amount was quantified (Pierce TM BCA Protein Assay Kit, Thermo Fisher Scientific, USA) and samples were diluted with loading buffer (10% sodium dodecyl sulfate, 312.5 mM Tris-HCl pH:6.8, 50% glycerol, and 0.05% bromophenol blue). Protein samples were subjected to SDS-PAGE (15 μg per lane) and blotted onto Protran Nitrocellulose Mem-brane-Whatman (GE Healthcare Life Science, USA). Membranes were blocked with Blocking Solution (5% powdered milk in TBST: Tris-buffered saline with 0.05% Tween-20) and incubated overnight at 4 °C with the respective anti-TRPM8 primary antibodies (see Table 2). Subsequently, membranes were washed with TBST, incubated with horseradish peroxidase (HRP) conjugated secondary antibody and developed with ECLplus (Thermo Fisher Scientific, USA) and imaged in an Amersham Imager 680 device. After several washes with TBST, membranes were incubated overnight at 4°C with anti-GFP (1:1000) and anti-GAPDH (1:5000) antibodies, developed and im-aged as explained before.

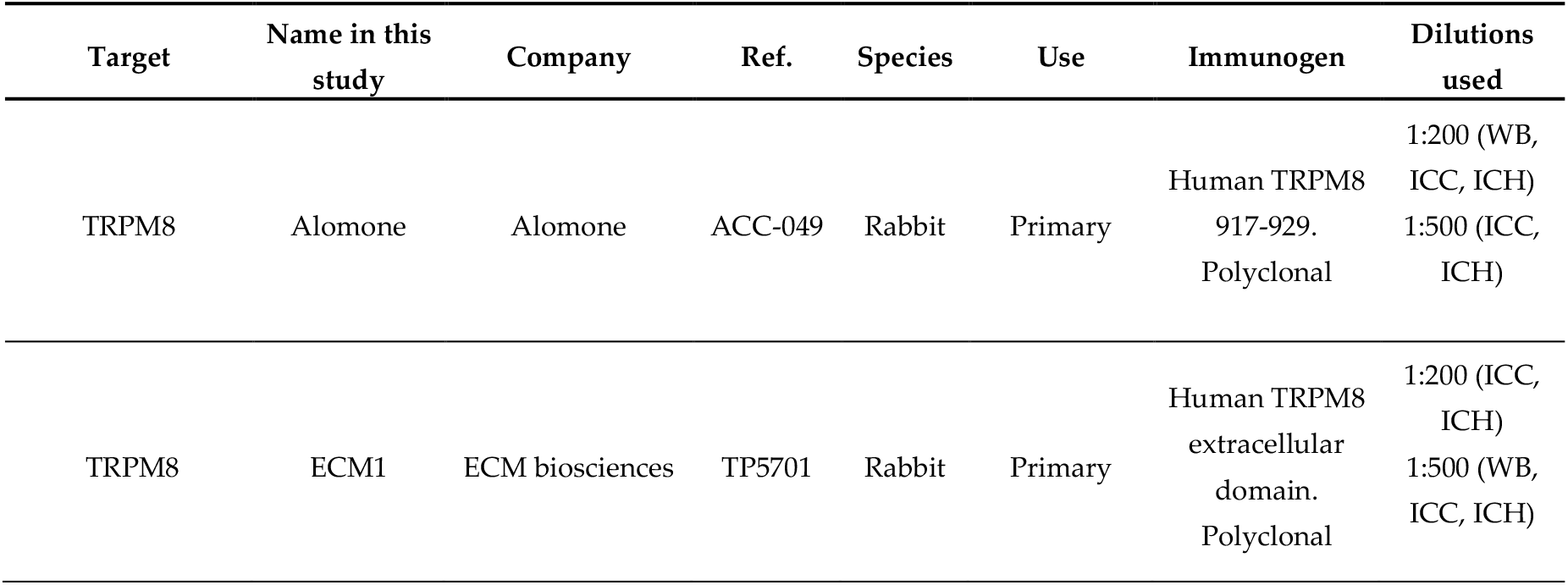
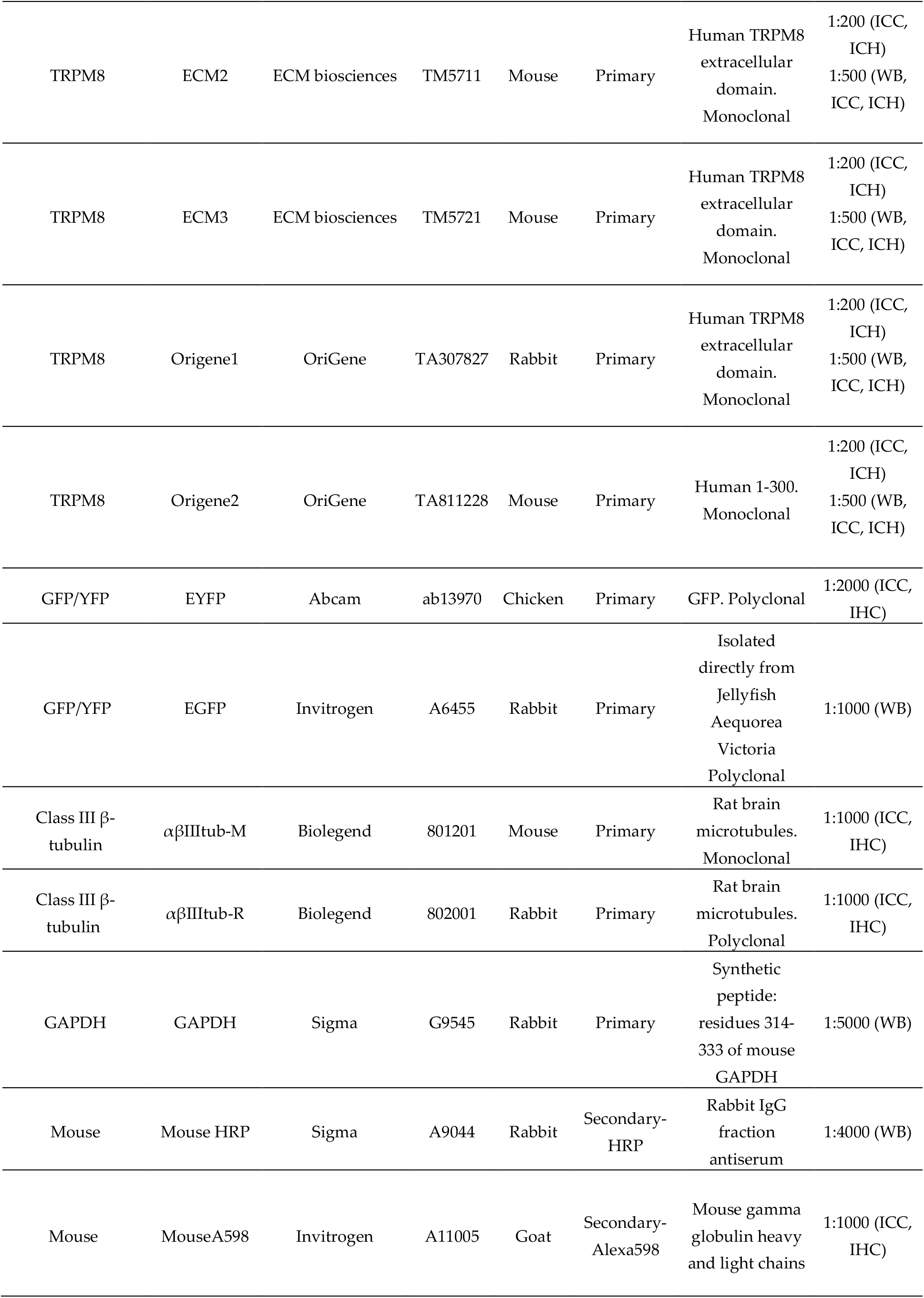
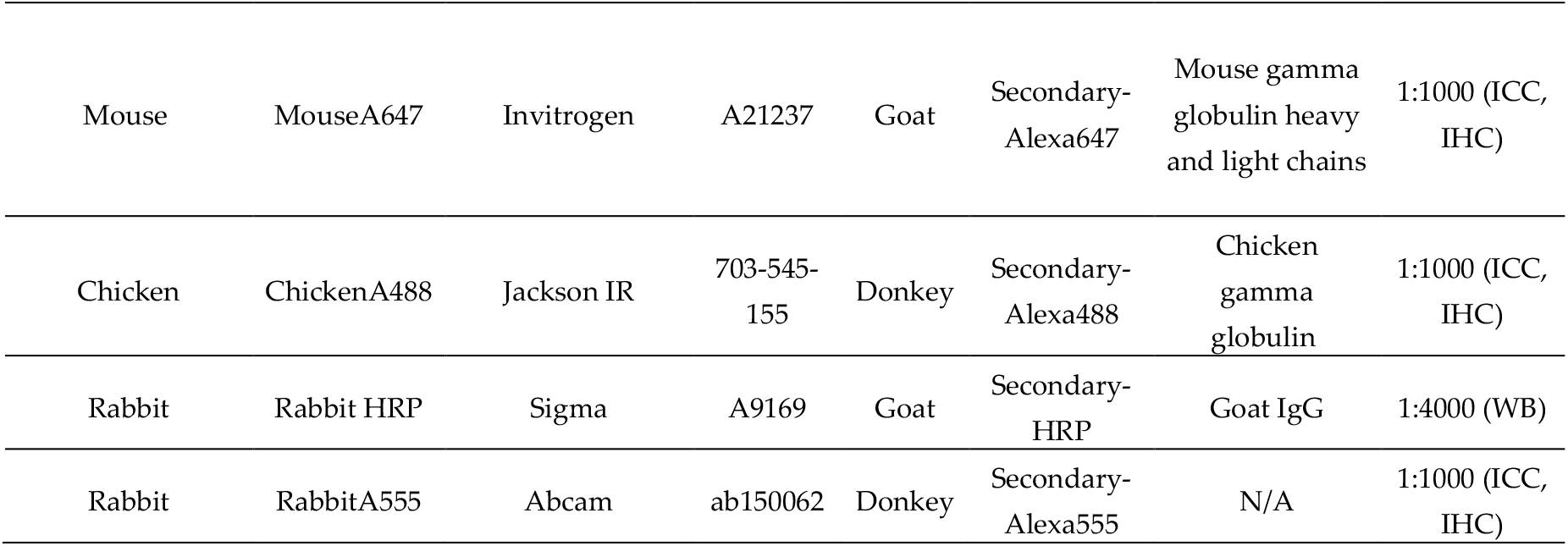

### Immunocytochemistry

Cells, cultured on coverslips, were fixed 10 min in 4 % PFA in 0.1M PBS, washed three times in PBS and twice in TTBS (0.5 M Tris Base, 9% w/v NaCl, 0.5% Tween 20, pH 7.6) 10 min. Non-specific binding sites were blocked by incubating the cells for 30 min in freshly-prepared blocking solution (1X TTBS, 1% bovine serum albumin (Tocris Bioscience, UK) and 0.25% Triton-X100). Cells were then incubated for 2 h at RT with the primary antibodies. Afterwards, cells were washed three times with TTBS 10 min and incubated for 45 min at RT with the secondary antibodies. Both primary and secondary antibodies were diluted in blocking solution. Cells were washed again three times with TTBS. An extra step was added for HEK cells, that were incubated for 5 min in Hoechst 33342 (Invitrogen, USA). Then, cells were washed with PBS and once with ddH2O and mounted on a microscope slide using VectaShield H-1000 antifade mounting medium (Vector Laboratories, USA). Mounted coverslips were sealed with clear nail polish and stored at 4°C until imaging. Z-stack images with a step size of ∼ 2 μm were acquired with an UPlanSApo 20× or a PlanApoN 60x objective using an inverted confocal microscope (FV1200, Olympus) driven by FV10-ASW 4.2 software (Olympus Life Sciences).

### Immunohistochemistry

Whole DRGs were fixed for 1 hour in 4% PFA in 0.1 M PBS, washed twice in PBS 10 min and cryoprotected in 30% sucrose overnight at 4 °C. Next day, tissue was embedded in optimal cutting temperature compound (OCT, Tissue-Tek) and frozen in dry ice. 20 μm slices were cut using an MNT cryostat (Slee Medical, Germany) and placed on SuperFrost microscope slides (Thermo Fisher Scientific, USA). Slides were dried in an oven at 37 °C for 30 min and washed twice with PBT (0.1 M PB, 0.05% Tween20, pH 7.4). Antigen retrieval (when used) was performed by boiling the samples in pH 6.0 citrate buffer for 20 min using a water bath and letting them cool down in the bath for 10 minutes. Non-specific binding sites were blocked for 1 hour with a blocking solution containing 5% bovine serum albumin and 1% Triton X-100 in PBT. Slides were incu-bated with the primary antibodies overnight at 4 °C. Next day, slides were washed four times with PBT 10 min each and incubated with the secondary antibodies for 2h at RT. Both primary and secondary antibodies were diluted as shown in Table 2 in block-ing solution. Then, slides were washed four times with PBT 10 min, once with PBS 10 min and once with ddH2O 5 min. Finally, slides were dried in dark conditions at RT and a glass coverslip was mounted on top of each slide using Fluoromount mounting medium (Sigma Aldrich, USA). Z-stack images with a step size of ∼ 2 μm were acquired with an UPlanSApo 20× or a PlanApoN 60x objective using an inverted confo-cal microscope (FV1200, Olympus) driven by FV10-ASW 4.2 software (Olympus Life Sciences).

### Image analysis and quantification

All quantitative measurements were performed using the 16-bit raw maximum projection images without any further modification. We defined the antibody specific-ity ratio (SR) in a TRPM8+ cell as the mean signal intensity of the antibody fluorescence in that cell divided by the average of the mean signal intensity in several (at least 20) TRMP8-cells in the same field. A SR value of 1 indicates the same antibody fluorescence intensity in TRPM8+ and TRPM8-cells, while SR > 1 indicates higher fluorescence intensity in TRPM8+ than in TRPM8-cells. TRPM8+ cells were identified by EYFP immunofluorescence, either in transfected cells or using a reporter mouse line. Images were analyzed with ImageJ 1.51j8 (NIH, USA) and Origin 2019 (OriginLab, USA).

### Image display

Immunofluorescence image brightness and contrast were only adjusted for image presentation to enhance visualization without altering any other feature. In the case of the TRPM8 antibody signal images, the maximum red pseudocolor level was set for the maximum fluorescence intensity, thus images displaying high background noise cor-respond to experiments in which the antibody signal was poor.

### Statistical analysis

Values are given as indicated in each figure caption. Normality of data distribution was checked with Shapiro-Wilk and Kolmogorov-Smirnov tests. Statistical significance was estimated with the Mann-Whitney test or Kruskal-Wallis followed by a Dunn’s posthoc test. A p-value < 0.05 was considered statistically significant. Analysis was performed using Prism version 8 (GraphPad Software, USA).

### Antibodies used in this study

All primary and secondary antibodies used are summarized in Table 2

## Supplementary Materials

Figure S1: Subcellular distribution of TRPM8 immunostaining in mTRPM8-EYFP.

Figure S2: Control immunostainings without primary antibodies.

Figure S3: Comparison of TRPM8 immunostaining in the TRPM8 reporter mouse compared to TRPM8KO mouse and to control without primary antibodies. Figure S4: Uncropped western blot membranes.

## Author Contributions

Conceptualization, JFT, EP and PHO.; methodology, PHO, RTM, EP and JFT; validation, JFT; formal analysis, PHO and JFT.; investigation, PHO, RTM, EP and JFT; re-sources, EP and FV; data curation, PHO, EP and JFT.; writing—original draft preparation, JFT.; writing—review and editing, JFT, FV, PHO and EP.; visualization, PHO.; supervision, JFT, EP and FV.; project administration, JFT; funding acquisition, FV and EP. All authors have read and agreed to the published version of the manuscript.

## Funding

During the course of this work, PH was supported by a FPI predoctoral fellowship (BES-2017-080782). This study was funded by MICIN/AEI/10.13039/501100011033 (PID2019-108194RB-I00), Generalitat Valenciana (PROMETEO/2021/031) and the Severo Ochoa Programme for Centres of Excellence in R&D (ref. SEV-2017-0723).

## Institutional Review Board Statement

All experimental procedures abide to the Spanish Royal Decree 53/2013 and the European Community Council directive 2016/63/EU, regulating the use of animals in research. The Committee on Animal Research at the University Miguel Hernández approved all the animal procedures.

## Data Availability Statement

Data available from the authors on reasonable request.

## Acknowledgments

We thank Ana Gomis for general support and supply of various reagents, antibodies and lab equipment. We thank all members of the Sensory Transduction and Nociception Group (www.painchannels.com) for insightful discussions. We thank Ardem Patapoutian and Ajay Dhaka (Scripps Research Institute and University of Washington) for providing Trpm8EGFPf; B6;129S1(FVB)-Trpm8tm1Apat/J knockout mouse line. The authors are grateful to Eva Quintero for excellent technical assistance and to Cristina Cahiz and Pablo Llorca for helping with some experiments.

## Conflicts of Interest

The authors declare no conflict of interest.

## Notes

### Competing Interest Statement

The authors have declared no competing interest.

## References

1. Peier, A.M.; Moqrich, A.; Hergarden, A.C.; Reeve, A.J.; Andersson, D.A.; Story, G.M.; Earley, T.J.; Dragoni, I.; McIntyre, P.; Bevan, S.; et al. A TRP Channel That Senses Cold Stimuli and Menthol. Cell 2002, 108, 705–715, doi:10.1016/s0092-8674(02)00652-9.

2. Voets, T.; Owsianik, G.; Nilius, B. TRPM8. In Transient Receptor Potential (TRP) Channels; Flockerzi, V., Nilius, B., Eds.; Handbook of Experimental Pharmacology; Springer Berlin Heidelberg: Berlin, Heidelberg, 2007; Vol. 179, pp. 329–344 ISBN 978-3-540-34889-4.

3. Ma, S.; G, G.; Ak, V.-E. Jf D. H H. Menthol Derivative WS-12 Selectively Activates Transient Receptor Potential Melastatin-8 (TRPM8) Ion Channels. Pak. J. Pharm. Sci. 2008, 21, 370–378.

4. Almaraz, L.; Manenschijn, J.-A.; de la Peña, E.; Viana, F. TRPM8. Handb. Exp. Pharmacol. 2014, 222, 547–579, doi:10.1007/978-3-642-54215-2_22.

5. McKemy, D.D.; Neuhausser, W.M.; Julius, D. Identification of a Cold Receptor Reveals a General Role for TRP Channels in Thermosensation. Nature 2002, 416, 52–58, doi:10.1038/nature719.

6. Qin, F. Regulation of TRP Ion Channels by Phosphatidylinositol-4,5-Bisphosphate. Handb. Exp. Pharmacol. 2007, 509–525, doi:10.1007/978-3-540-34891-7_30.

7. Rohács, T.; Lopes, C.M.B.; Michailidis, I.; Logothetis, D.E. PI(4,5)P2 Regulates the Activation and Desensitization of TRPM8 Channels through the TRP Domain. Nat. Neurosci. 2005, 8, 626–634, doi:10.1038/nn1451.

8. Vriens, J.; Nilius, B.; Voets, T. Peripheral Thermosensation in Mammals. Nat. Rev. Neurosci. 2014, 15, 573–589, doi:10.1038/nrn3784.

9. Parra, A.; Madrid, R.; Echevarria, D.; del Olmo, S.; Morenilla-Palao, C.; Acosta, M.C.; Gallar, J.; Dhaka, A.; Viana, F.; Belmonte, C. Ocular Surface Wetness Is Regulated by TRPM8-Dependent Cold Thermoreceptors of the Cornea. Nat. Med. 2010, 16, 1396–1399, doi:10.1038/nm.2264.

10. Robbins, A.; Kurose, M.; Winterson, B.J.; Meng, I.D. Menthol Activation of Corneal Cool Cells Induces TRPM8-Mediated Lacrimation but Not Nociceptive Responses in Rodents. Investig. Opthalmology Vis. Sci. 2012, 53, 7034, doi:10.1167/iovs.12-10025.

11. Quallo, T.; Vastani, N.; Horridge, E.; Gentry, C.; Parra, A.; Moss, S.; Viana, F.; Belmonte, C.; Andersson, D.A.; Bevan, S. TRPM8 Is a Neuronal Osmosensor That Regulates Eye Blinking in Mice. Nat. Commun. 2015, 6, 7150, doi:10.1038/ncomms8150.

12. Belmonte, C.; Gallar, J. Cold Thermoreceptors, Unexpected Players in Tear Production and Ocular Dryness Sensations. Investig. Opthalmology Vis. Sci. 2011, 52, 3888, doi:10.1167/iovs.09-5119.

13. Belmonte, C.; Nichols, J.J.; Cox, S.M.; Brock, J.A.; Begley, C.G.; Bereiter, D.A.; Dartt, D.A.; Galor, A.; Hamrah, P.; Ivanusic, J.J.; et al. TFOS DEWS II Pain and Sensation Report. Ocul. Surf. 2017, 15, 404–437, doi:10.1016/j.jtos.2017.05.002.

14. Ma, S.; Yu, H.; Zhao, Z.; Luo, Z.; Chen, J.; Ni, Y.; Jin, R.; Ma, L.; Wang, P.; Zhu, Z.; et al. Activation of the Cold-Sensing TRPM8 Channel Triggers UCP1-Dependent Thermogenesis and Prevents Obesity. J. Mol. Cell Biol. 2012, 4, 88–96, doi:10.1093/jmcb/mjs001.

15. Henström, M.; Hadizadeh, F.; Beyder, A.; Bonfiglio, F.; Zheng, T.; Assadi, G.; Rafter, J.; Bujanda, L.; Agreus, L.; Andreasson, A.; et al. TRPM8 Polymorphisms Associated with Increased Risk of IBS-C and IBS-M. Gut 2017, 66, 1725–1727, doi:10.1136/gutjnl-2016-313346.

16. Hosoya, T.; Matsumoto, K.; Tashima, K.; Nakamura, H.; Fujino, H.; Murayama, T.; Horie, S. TRPM8 Has a Key Role in Experimental Colitis-Induced Visceral Hyperalgesia in Mice. Neurogastroenterol. Motil. Off. J. Eur. Gastrointest. Motil. Soc. 2014, 26, 1112–1121, doi:10.1111/nmo.12368.

17. Ramachandran, R.; Hyun, E.; Zhao, L.; Lapointe, T.K.; Chapman, K.; Hirota, C.L.; Ghosh, S.; McKemy, D.D.; Vergnolle, N.; Beck, P.L.; et al. TRPM8 Activation Attenuates Inflammatory Responses in Mouse Models of Colitis. Proc. Natl. Acad. Sci. U. S. A. 2013, 110, 7476–7481, doi:10.1073/pnas.1217431110.

18. Gardiner, J.C.; Kirkup, A.J.; Curry, J.; Humphreys, S.; O’Regan, P.; Postlethwaite, M.; Young, K.C.; Kitching, L.; Ethell, B.T.; Winpenny, D.; et al. The Role of TRPM8 in the Guinea-Pig Bladder-Cooling Reflex Investigated Using a Novel TRPM8 Antagonist. Eur. J. Pharmacol. 2014, 740, 398–409, doi:10.1016/j.ejphar.2014.07.022.

19. Uvin, P.; Franken, J.; Pinto, S.; Rietjens, R.; Grammet, L.; Deruyver, Y.; Alpizar, Y.A.; Talavera, K.; Vennekens, R.; Everaerts, W.; et al. Essential Role of Transient Receptor Potential M8 (TRPM8) in a Model of Acute Cold-Induced Urinary Urgency. Eur. Urol. 2015, 68, 655–661, doi:10.1016/j.eururo.2015.03.037.

20. Andersson, K.-E.; Gratzke, C.; Hedlund, P. The Role of the Transient Receptor Potential (TRP) Superfamily of Cation-Selective Channels in the Management of the Overactive Bladder. BJU Int. 2010, 106, 1114–1127, doi:10.1111/j.1464-410X.2010.09650.x.

21. Mukerji, G.; Yiangou, Y.; Corcoran, S.L.; Selmer, I.S.; Smith, G.D.; Benham, C.D.; Bountra, C.; Agarwal, S.K.; Anand, P. Cool and Menthol Receptor TRPM8 in Human Urinary Bladder Disorders and Clinical Correlations. BMC Urol. 2006, 6, 6, doi:10.1186/1471-2490-6-6.

22. Ordás, P.; Hernández-Ortego, P.; Vara, H.; Fernández-Peña, C.; Reimúndez, A.; Morenilla-Palao, C.; Guadaño-Ferraz, A.; Gomis, A.; Hoon, M.; Viana, F.; et al. Expression of the Cold Thermoreceptor TRPM8 in Rodent Brain Thermoregulatory Circuits. J. Comp. Neurol. 2021, 529, 234–256, doi:10.1002/cne.24694.

23. Reimúndez, A.; Fernández-Peña, C.; Ordás, P.; Hernández-Ortego, P.; Gallego, R.; Morenilla-Palao, C.; Navarro, J.; Martín-Cora, F.; Pardo-Vázquez, J.L.; Schwarz, L.A.; et al. The Cold-Sensing Ion Channel TRPM8 Regulates Central and Peripheral Clockwork and the Circadian Oscillations of Body Temperature. Acta Physiol. n/a, e13896, doi:10.1111/apha.13896.

24. Valero, M.; Morenilla-Palao, C.; Belmonte, C.; Viana, F. Pharmacological and Functional Properties of TRPM8 Channels in Prostate Tumor Cells. Pflugers Arch. 2011, 461, 99–114, doi:10.1007/s00424-010-0895-0.

25. Dhaka, A.; Murray, A.N.; Mathur, J.; Earley, T.J.; Petrus, M.J.; Patapoutian, A. TRPM8 Is Required for Cold Sensation in Mice. Neuron 2007, 54, 371–378, doi:10.1016/j.neuron.2007.02.024.

26. takashima, Y.; Daniels, R.L.; Knowlton, W.; Teng, J.; Liman, E.R.; McKemy, D.D. Diversity in the Neural Circuitry of Cold Sensing Revealed by Genetic Axonal Labeling of Transient Receptor Potential Melastatin 8 Neurons. J. Neurosci. Off. J. Soc. Neurosci. 2007, 27, 14147–14157, doi:10.1523/JNEUROSCI.4578-07.2007.

27. Kobayashi, K.; Fukuoka, T.; Obata, K.; Yamanaka, H.; Dai, Y.; Tokunaga, A.; Noguchi, K. Distinct Expression of TRPM8, TRPA1, and TRPV1 MRNAs in Rat Primary Afferent Neurons with Aδ/c-Fibers and Colocalization with Trk Receptors. J. Comp. Neurol. 2005, 493, 596–606, doi:10.1002/cne.20794.

28. Meissner, M.; Obmann, V.C.; Hoschke, M.; Link, S.; Jung, M.; Held, G.; Philipp, S.E.; Zimmermann, R.; Flockerzi, V. Lessons of Studying TRP Channels with Antibodies. In TRP Channels; Zhu, M.X., Ed.; CRC Press/Taylor & Francis: Boca Raton (FL), 2011 ISBN 978-1-4398-1860-2.

29. Jemal, I.; Soriano, S.; Conte, A.L.; Morenilla, C.; Gomis, A. G Protein-Coupled Receptor Signalling Potentiates the Osmo-Mechanical Activation of TRPC5 Channels. Pflugers Arch. 2014, 466, 1635–1646, doi:10.1007/s00424-013-1392-z.

30. Veliz, L.A.; Toro, C.A.; Vivar, J.P.; Arias, L.A.; Villegas, J.; Castro, M.A.; Brauchi, S. Near-Membrane Dynamics and Capture of TRPM8 Channels within Transient Confinement Domains. PLoS ONE 2010, 5, doi:10.1371/journal.pone.0013290.

31. Ghosh, D.; Pinto, S.; Danglot, L.; Vandewauw, I.; Segal, A.; Van Ranst, N.; Benoit, M.; Janssens, A.; Vennekens, R.; Vanden Berghe, P.; et al. VAMP7 Regulates Constitutive Membrane Incorporation of the Cold-Activated Channel TRPM8. Nat. Commun. 2016, 7, 10489, doi:10.1038/ncomms10489.

32. Rivera, B.; Moreno, C.; Lavanderos, B.; Hwang, J.Y.; Fernández-Trillo, J.; Park, K.S.; Orio, P.; Viana, F.; Madrid, R.; Pertusa, M. Constitutive Phosphorylation as a Key Regulator of Trpm8 Channel Function. J. Neurosci. 2021, 41, 8475–8493, doi:10.1523/JNEUROSCI.0345-21.2021.

33. Morenilla-Palao, C.; Luis, E.; Fernández-Peña, C.; Quintero, E.; Weaver, J.L.; Bayliss, D.A.; Viana, F. Ion Channel Profile of TRPM8 Cold Receptors Reveals a Role of TASK-3 Potassium Channels in Thermosensation. Cell Rep. 2014, 8, 1571–1582, doi:10.1016/j.celrep.2014.08.003.

34. Cattoretti, G.; Pileri, S.; Parravicini, C.; Becker, M.H.; Poggi, S.; Bifulco, C.; Key, G.; D’Amato, L.; Sabattini, E.; Feudale, E. Antigen Unmasking on Formalin-Fixed, Paraffin-Embedded Tissue Sections. J. Pathol. 1993, 171, 83–98, doi:10.1002/path.1711710205.

35. Krenacs, L.; Krenacs, T.; Stelkovics, E.; Raffeld, M. Heat-Induced Antigen Retrieval for Immunohistochemical Reactions in Routinely Processed Paraffin Sections. Methods Mol. Biol. Clifton NJ 2010, 588, 103–119, doi:10.1007/978-1-59745-324-0_14.

36. Bordeaux, J.; Welsh, A.W.; Agarwal, S.; Killiam, E.; Baquero, M.T.; Hanna, J.A.; Anagnostou, V.K.; Rimm, D.L. Antibody Validation. BioTechniques 2010, 48, 197–209, doi:10.2144/000113382.

37. Burry, R.W. Controls for Immunocytochemistry: An Update. J. Histochem. Cytochem. 2011, 59, 6–12, doi:10.1369/jhc.2010.956920.

38. Möller, M.; Möser, C.V.; Weiß, U.; Niederberger, E. The Role of AlphαSynuclein in Mouse Models of Acute, Inflammatory and Neuropathic Pain. Cells 2022, 11, 1967, doi:10.3390/cells11121967.

39. Genovesi, S.; Moro, R.; Vignoli, B.; De Felice, D.; Canossa, M.; Montironi, R.; Carbone, F.G.; Barbareschi, M.; Lunardi, A.; Alaimo, A. Trpm8 Expression in Human and Mouse Castration Resistant Prostate Adenocarcinoma Paves the Way for the Preclinical Development of TRPM8-Based Targeted Therapies. Biomolecules 2022, 12, 193, doi:10.3390/biom12020193.

40. thapa, D.; Barrett, B.; Argunhan, F.; Brain, S.D. Influence of Cold-TRP Receptors on Cold-Influenced Behaviour. Pharmaceuticals 2021, 15, 42, doi:10.3390/ph15010042.

41. Vinuela-Fernandez, I.; Sun, L.; Jerina, H.; Curtis, J.; Allchorne, A.; Gooding, H.; Rosie, R.; Holland, P.; Tas, B.; Mitchell, R.; et al. The TRPM8 Channel Forms a Complex with the 5-HT1B Receptor and Phospholipase D That Amplifies Its Reversal of Pain Hypersensitivity. Neuropharmacology 2014, 79, 136–151, doi:10.1016/j.neuropharm.2013.11.006.

42. Gkika, D.; Lemonnier, L.; Shapovalov, G.; Gordienko, D.; Poux, C.; Bernardini, M.; Bokhobza, A.; Bidaux, G.; Degerny, C.; Verreman, K.; et al. TRP Channel-Associated Factors Are a Novel Protein Family That Regulates TRPM8 Trafficking and Activity. J. Cell Biol. 2015, 208, 89–107, doi:10.1083/jcb.201402076.

43. Yudin, Y.; Lutz, B.; Tao, Y.-X.; Rohacs, T. Phospholipase C Δ4 Regulates Cold Sensitivity in Mice. J. Physiol. 2016, 594, 3609–3628, doi:10.1113/JP272321.

44. Chukyo, A.; Chiba, T.; Kambe, T.; Yamamoto, K.; Kawakami, K.; Taguchi, K.; Abe, K. Oxaliplatin-Induced Changes in Expression of Transient Receptor Potential Channels in the Dorsal Root Ganglion as a Neuropathic Mechanism for Cold Hypersensitivity. Neuropeptides 2018, 67, 95–101, doi:10.1016/j.npep.2017.12.002.

45. Villalba-Riquelme, E.; de la Torre-Martínez, R.; Fernández-Carvajal, A.; Ferrer-Montiel, A. Paclitaxel in Vitro Reversibly Sensitizes the Excitability of IB4(-) and IB4(+) Sensory Neurons from Male and Female Rats. Br. J. Pharmacol. 2022, 179, 3693–3710, doi:10.1111/bph.15809.

46. Acharya, T.K.; Kumar, S.; Tiwari, N.; Ghosh, A.; Tiwari, A.; Pal, S.; Majhi, R.K.; Kumar, A.; Das, R.; Singh, A.; et al. TRPM8 Channel Inhibitor-Encapsulated Hydrogel as a Tunable Surface for Bone Tissue Engineering. Sci. Rep. 2021, 11, 3730, doi:10.1038/s41598-021-81041-w.

47. Lee, P.R.; Lee, J.Y.; Kim, H.B.; Lee, J.H.; Oh, S.B. TRPM8 Mediates Hyperosmotic Stimuli-Induced Nociception in Dental Afferents. J. Dent. Res. 2020, 99, 107–114, doi:10.1177/0022034519886847.

48. Majhi, R.K.; Saha, S.; Kumar, A.; Ghosh, A.; Swain, N.; Goswami, L.; Mohapatra, P.; Maity, A.; Kumar Sahoo, V.; Kumar, A.; et al. Expression of Temperature-Sensitive Ion Channel TRPM8 in Sperm Cells Correlates with Vertebrate Evolution. PeerJ 2015, 3, e1310, doi:10.7717/peerj.1310.

49. Cornejo, V.H.; González, C.; Campos, M.; Vargas-Saturno, L.; Juricic, M. de los Á.; Miserey-Lenkei, S.; Pertusa, M.; Madrid, R.; Couve, A. Non-Conventional Axonal Organelles Control TRPM8 Ion Channel Trafficking and Peripheral Cold Sensing. Cell Rep. 2020, 30, 4505-4517.e5, doi:10.1016/j.celrep.2020.03.017.

50. He, J.; Pham, T.L.; Kakazu, A.H.; Bazan, H.E.P. Remodeling of Substance P Sensory Nerves and Transient Receptor Potential Melastatin 8 (TRPM8) Cold Receptors After Corneal Experimental Surgery. Investig. Opthalmology Vis. Sci. 2019, 60, 2449, doi:10.1167/iovs.18-26384.

51. Borowiec, A.; Sion, B.; Chalmel, F.; Rolland, A.D.; Lemonnier, L.; De Clerck, T.; Bokhobza, A.; Derouiche, S.; Dewailly, E.; Slomianny, C.; et al. Cold/Menthol TRPM8 Receptors Initiate the Cold-shock Response and Protect Germ Cells from Cold-shock–Induced Oxidation. FASEB J. 2016, 30, 3155–3170, doi:10.1096/fj.201600257R.

52. Ramos-Vara, J.A.; Miller, M.A. When Tissue Antigens and Antibodies Get Along: Revisiting the Technical Aspects of Immunohistochemistry—The Red, Brown, and Blue Technique. Vet. Pathol. 2014, 51, 42–87, doi:10.1177/0300985813505879.

53. Dhaka, A.; Earley, T.J.; Watson, J.; Patapoutian, A. Visualizing Cold Spots: TRPM8-Expressing Sensory Neurons and Their Projections. J. Neurosci. 2008, 28, 566–575, doi:10.1523/JNEUROSCI.3976-07.2008.

54. Pertusa, M.; González, A.; Hardy, P.; Madrid, R.; Viana, F. Bidirectional Modulation of Thermal and Chemical Sensitivity of TRPM8 Channels by the Initial Region of the N-Terminal Domain. J. Biol. Chem. 2014, 289, 21828–21843, doi:10.1074/jbc.M114.565994.

55. Caires, R.; Luis, E.; Taberner, F.J.; Fernandez-Ballester, G.; Ferrer-Montiel, A.; Balazs, E.A.; Gomis, A.; Belmonte, C.; De La Peña, E. Hyaluronan Modulates TRPV1 Channel Opening, Reducing Peripheral Nociceptor Activity and Pain. Nat. Commun. 2015, 6, doi:10.1038/ncomms9095.

