## Supplementary Figures for "Validation of six commercial antibodies for detection of heterologous and endogenous TRPM8 ion channel expression"

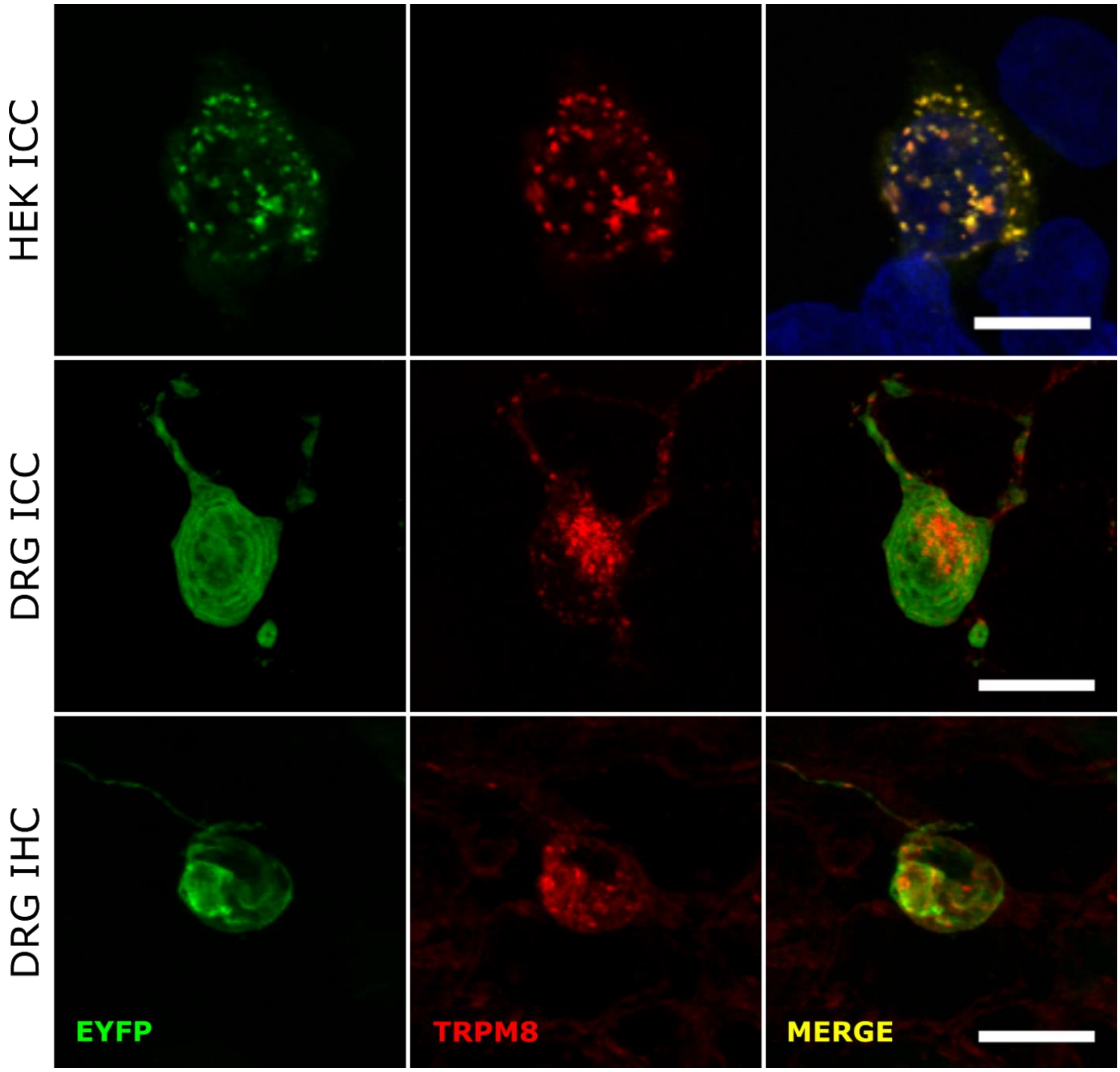


**FIGURE S1**. **Subcellular distribution of TRPM8 immunostaining in mTRPM8-EYFP transfected HEK cells ICC and DRG ICC and IHC.** Confocal images showing the fluorescent pattern of TRPM8 staining. Green: EYFP. Red: TRPM8 (Origene1 antibody). Blue: Hoechst. Scale bar: 10μm for HEK ICC, 15 μm for DRG ICC and IHC.

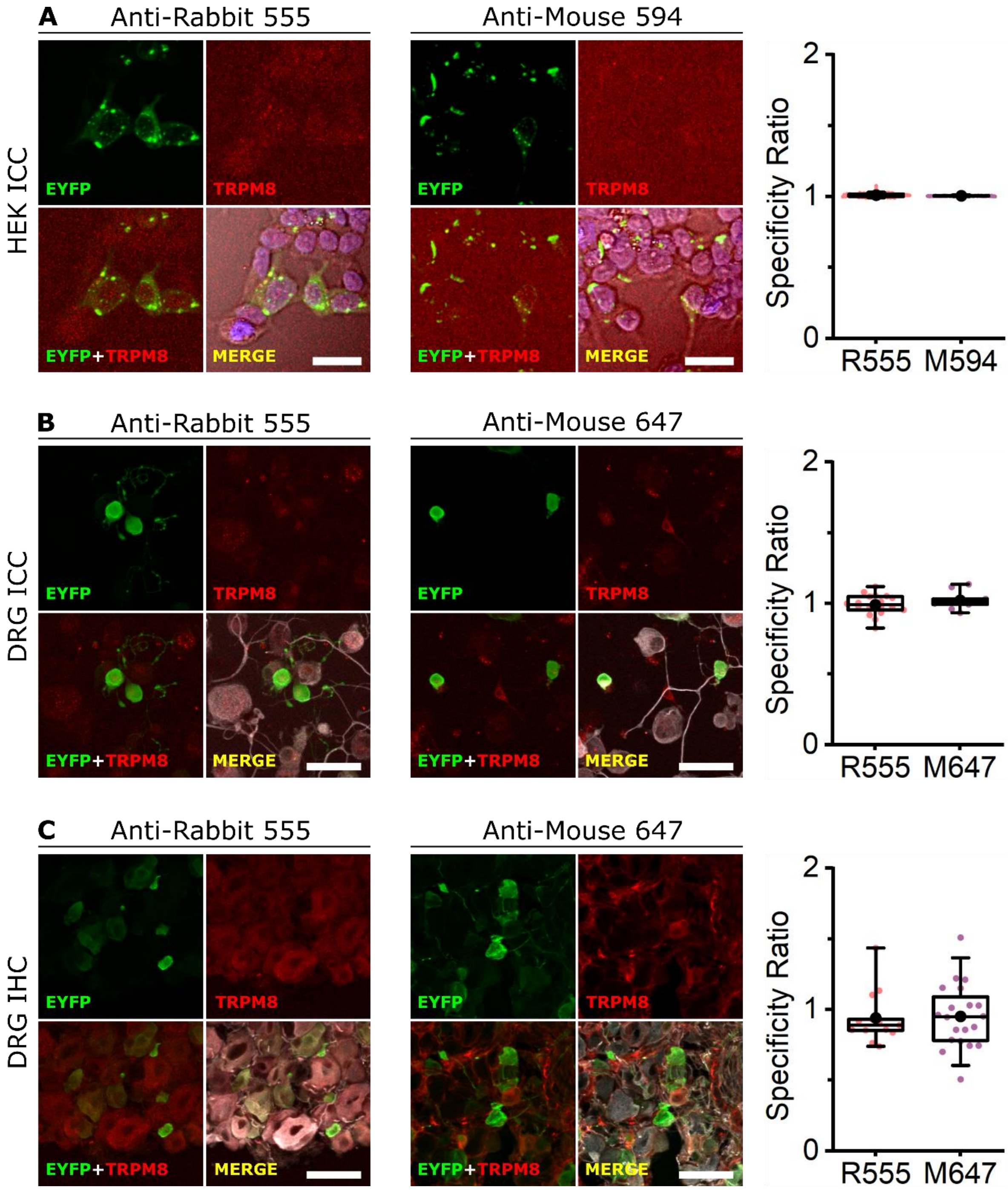


**FIGURE S2**. **Control immunostainings without primary antibodies. A, B, C** left, middle. Confocal images of immunochemistry in transfected HEK-293 cells (**A**), cultured DRG neurons (**B**) and DRG slices (**C**) from the *Trpm8$^{BAC}$-EYFP$^{+}$* mouse. EYFP (green), no TRPM8 primary antibody (red), Hoechst (blue), βIII-Tubulin (grey). Merge in A correspond to the overlap of the three fluorescent signals plus the bright field image. Scale bar: 30 µm for A, 50µm for B and C. **A,B,C** right**.** Box plots represent specificity ratio (SR) for each dilution of the respective antibody. Each dot corresponds to an individual EYFP+ cell. Box contains the 25$^{th}$ to 75$^{th}$ percentiles. Whiskers mark the 5$^{th}$ and 95$^{th}$ percentiles. The line inside the box denotes the median and the black dot represents the mean. n = at least 10 cells; 2 images from 1 mouse for each secondary antibody.

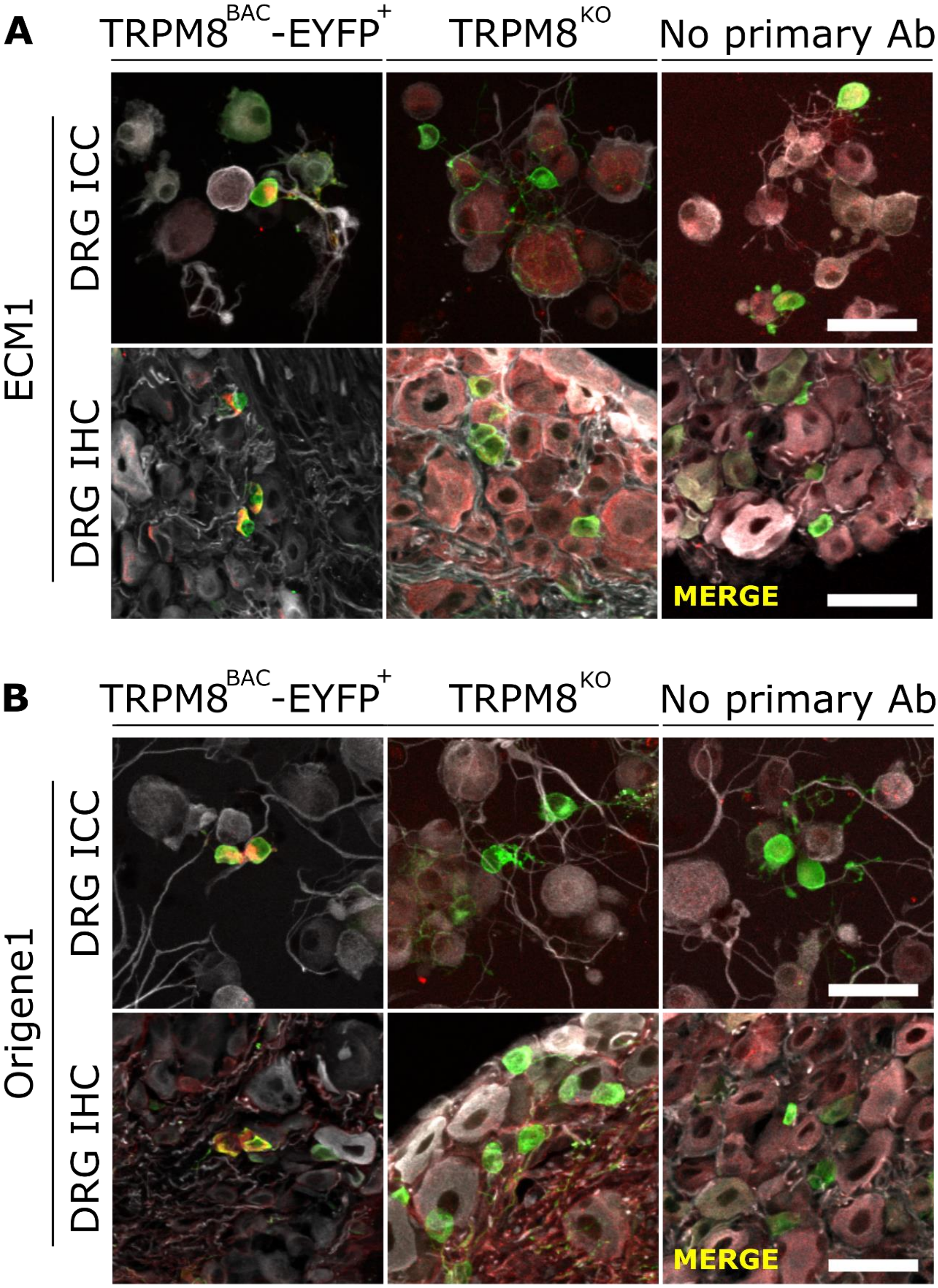


**FIGURE S3 Comparison of TRPM8 immunostaining in the TRPM8 reporter mouse compared to TRPM8$^{KO}$ mouse and to control without primary antibodies.** Confocal merged images of ICC and IHC in DRGs using the ECM1 (**A**) and the Origene1 (**B)** antibodies. EYFP (green) TRPM8/secondary antibody alone (red), βIII-tubulin (grey). Scale bar: 50 µm.

### ALOMONE

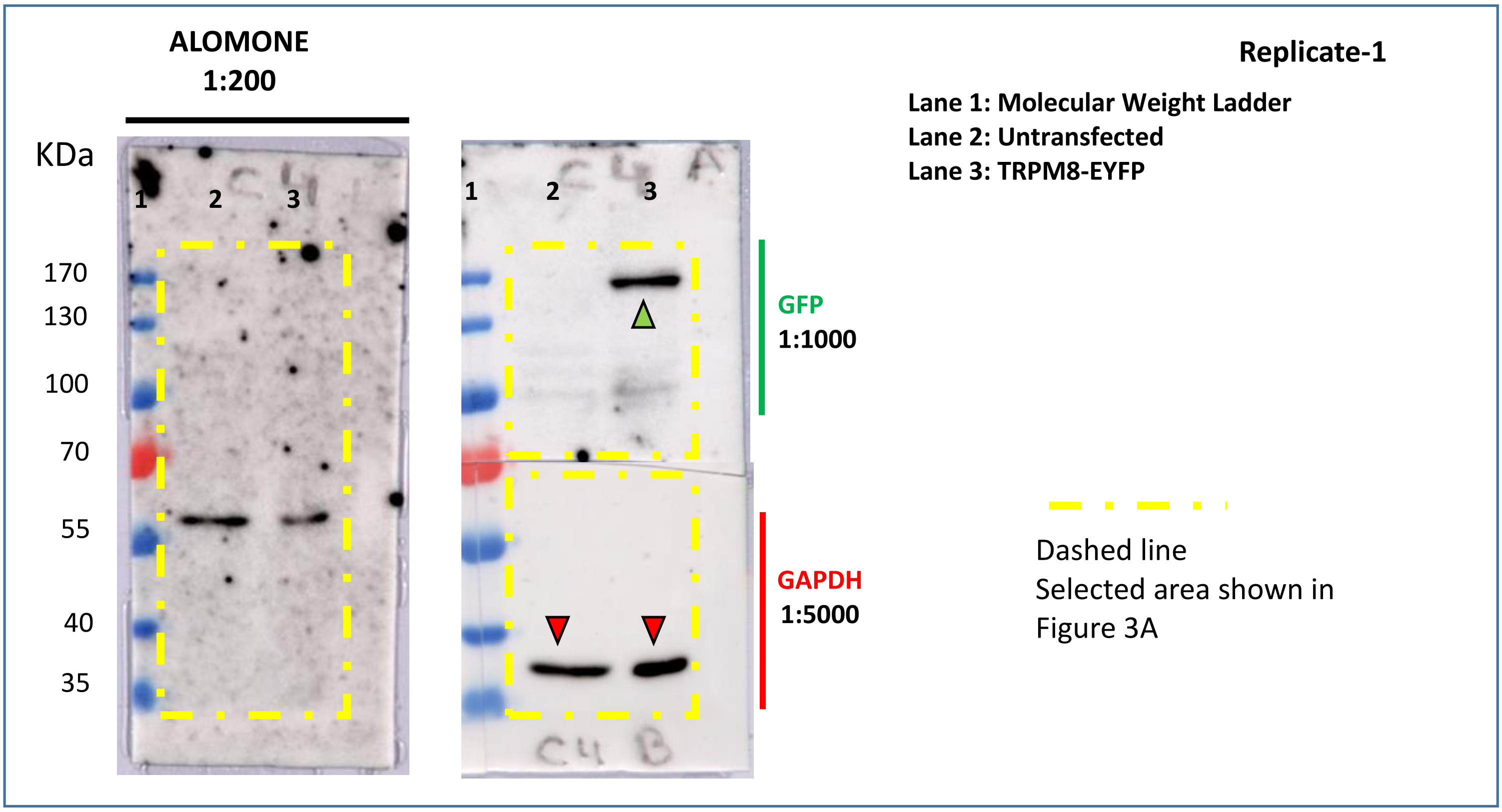

ALOMONE
1:200
KDa
170
130
100
70
55
40
35
1 2 3
1 2 3
GFP
1:1000
GAPDH
1:5000
Replicate-1
Lane 1: Molecular Weight Ladder
Lane 2: Untransfected
Lane 3: TRPM8-EYFP
Dashed line
Selected area shown in
Figure 3A


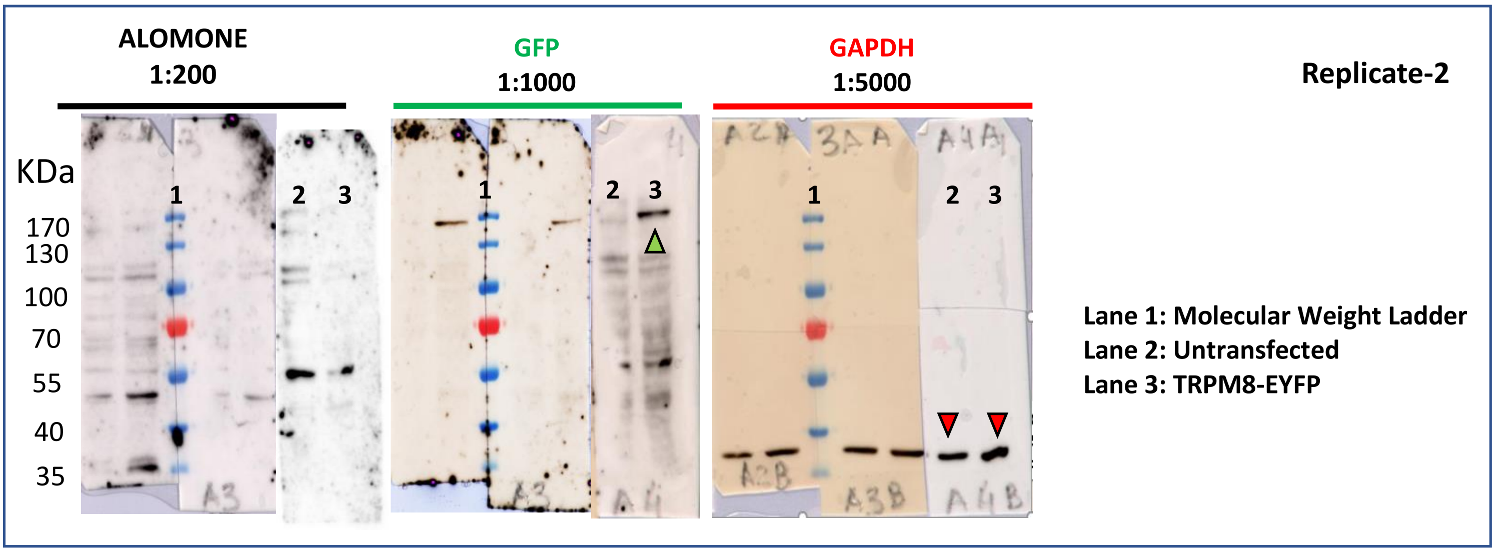

ALOMONE
1:200
GFP
1:1000
GAPDH
1:5000
Replicate-2
KDa
170
130
100
70
55
40
35
1 2 3
1 2 3
1 2 3
Lane 1: Molecular Weight Ladder
Lane 2: Untransfected
Lane 3: TRPM8-EYFP


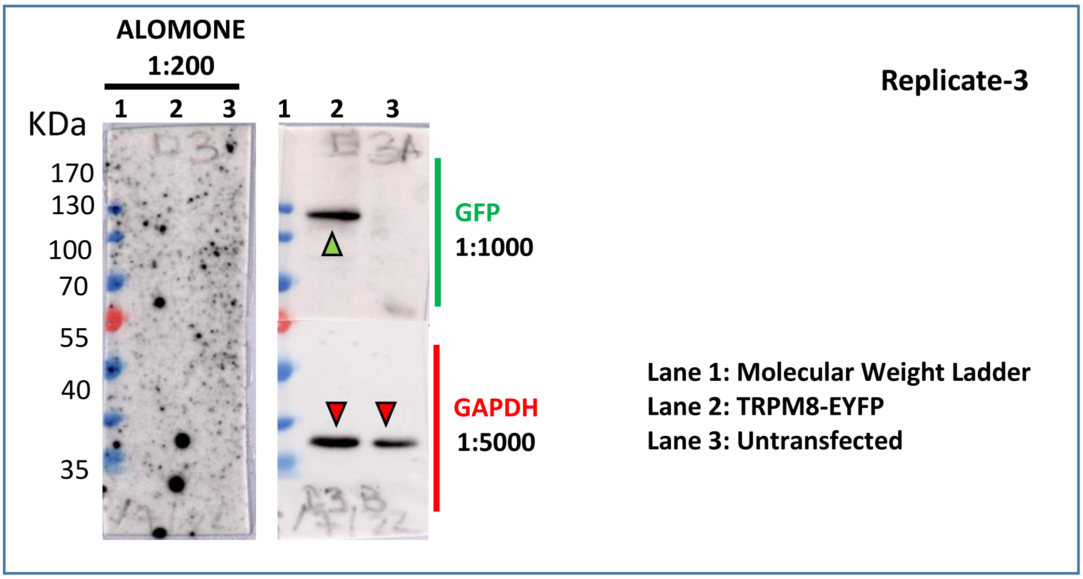

ALOMONE
1:200
Replicate-3
1 2 3
1 2 3
KDa
170
130
100
70
55
40
35
GFP
1:1000
GAPDH
1:5000
Lane 1: Molecular Weight Ladder
Lane 2: TRPM8-EYFP
Lane 3: Untransfected

### ECM1

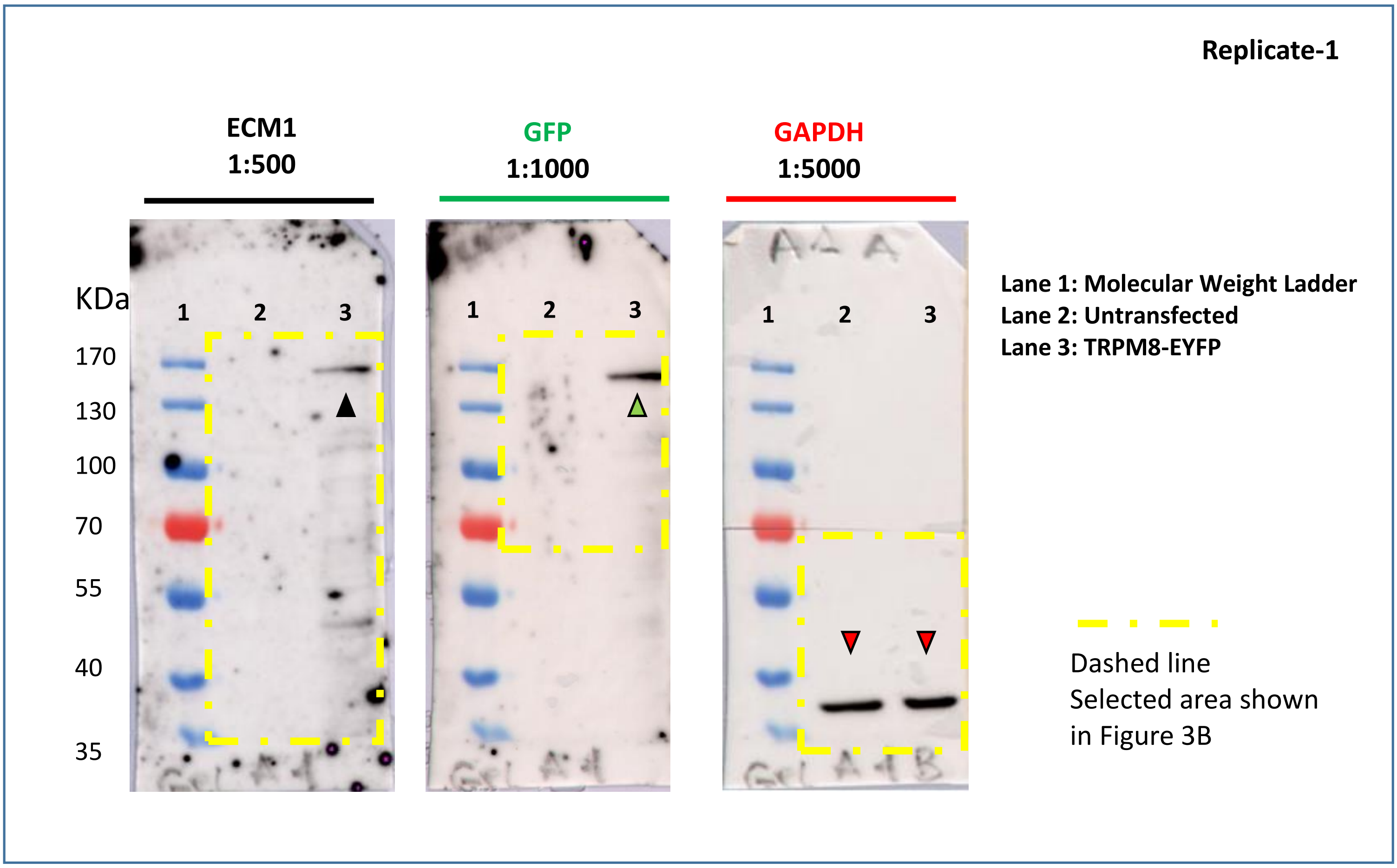

Replicate-1
ECM1
1:500
GFP
1:1000
GAPDH
1:5000
KDa
170
130
100
70
55
40
35
1
2
3
Lane 1: Molecular Weight Ladder
Lane 2: Untransfected
Lane 3: TRPM8-EYFP
Dashed line
Selected area shown
in Figure 3B

### ECM2

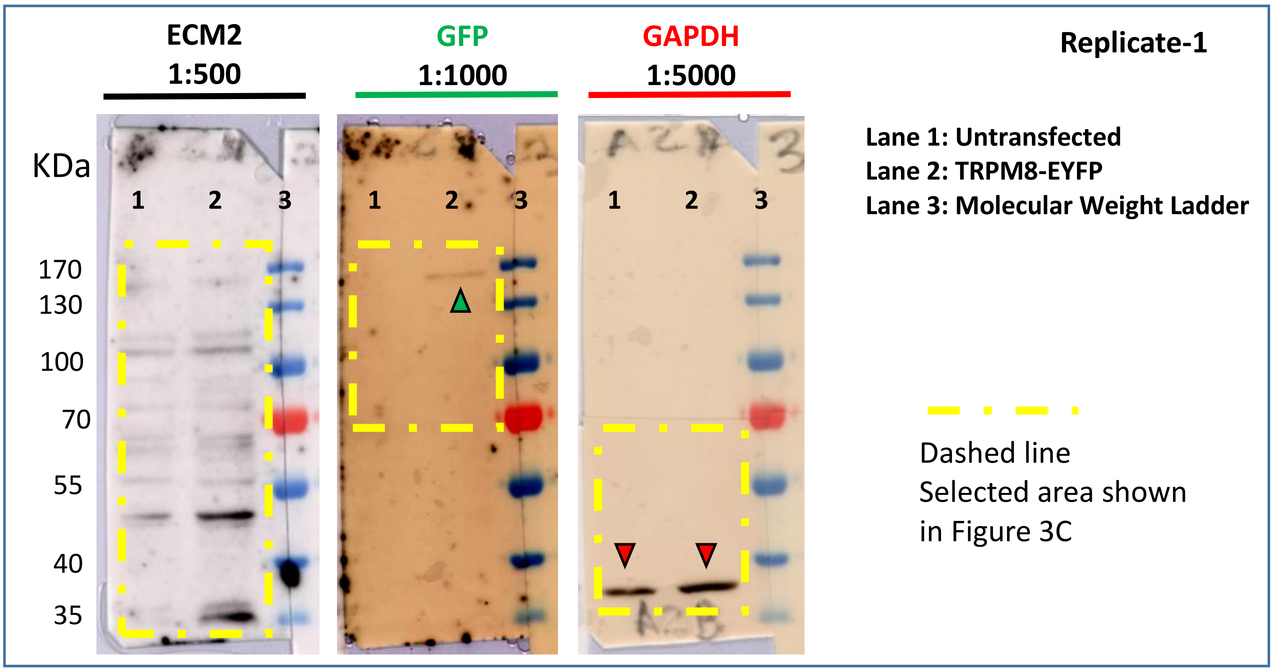

ECM2
1:500
GFP
1:1000
GAPDH
1:5000
Replicate-1
KDa
1 2 3
170
130
100
70
55
40
35
Lane 1: Untransfected
Lane 2: TRPM8-EYFP
Lane 3: Molecular Weight Ladder
Dashed line
Selected area shown
in Figure 3C


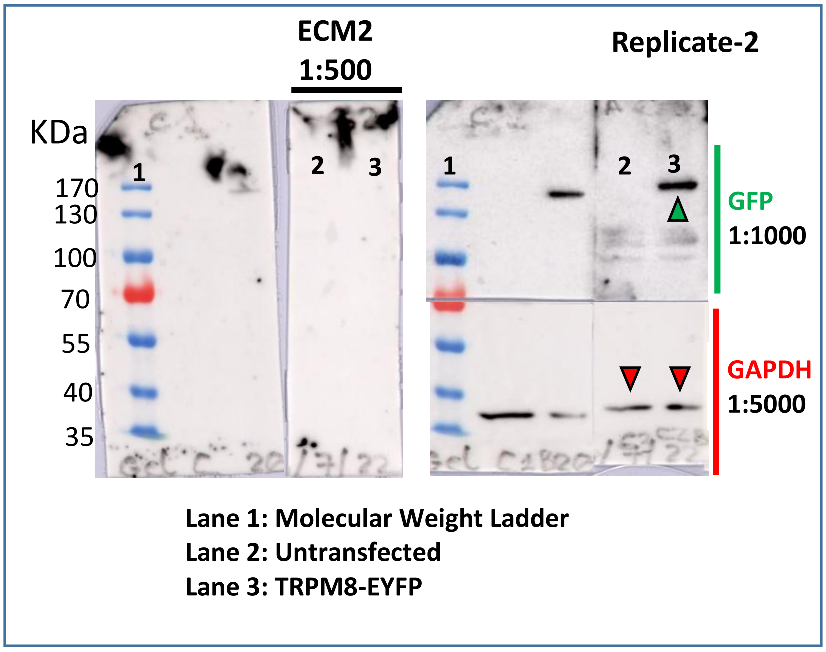

ECM2
1:500
Replicate-2
KDa
1 2 3
170
130
100
70
55
40
35
GFP
1:1000
GAPDH
1:5000
Lane 1: Molecular Weight Ladder
Lane 2: Untransfected
Lane 3: TRPM8-EYFP


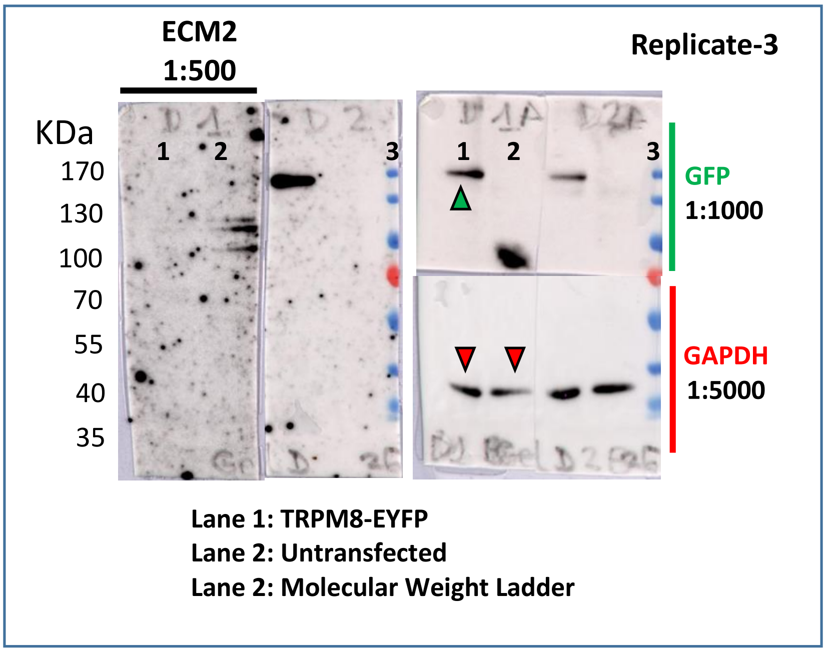

ECM2
1:500
Replicate-3
KDa
1 2 3
170
130
100
70
55
40
35
GFP
1:1000
GAPDH
1:5000
Lane 1: TRPM8-EYFP
Lane 2: Untransfected
Lane 2: Molecular Weight Ladder

### ECM3

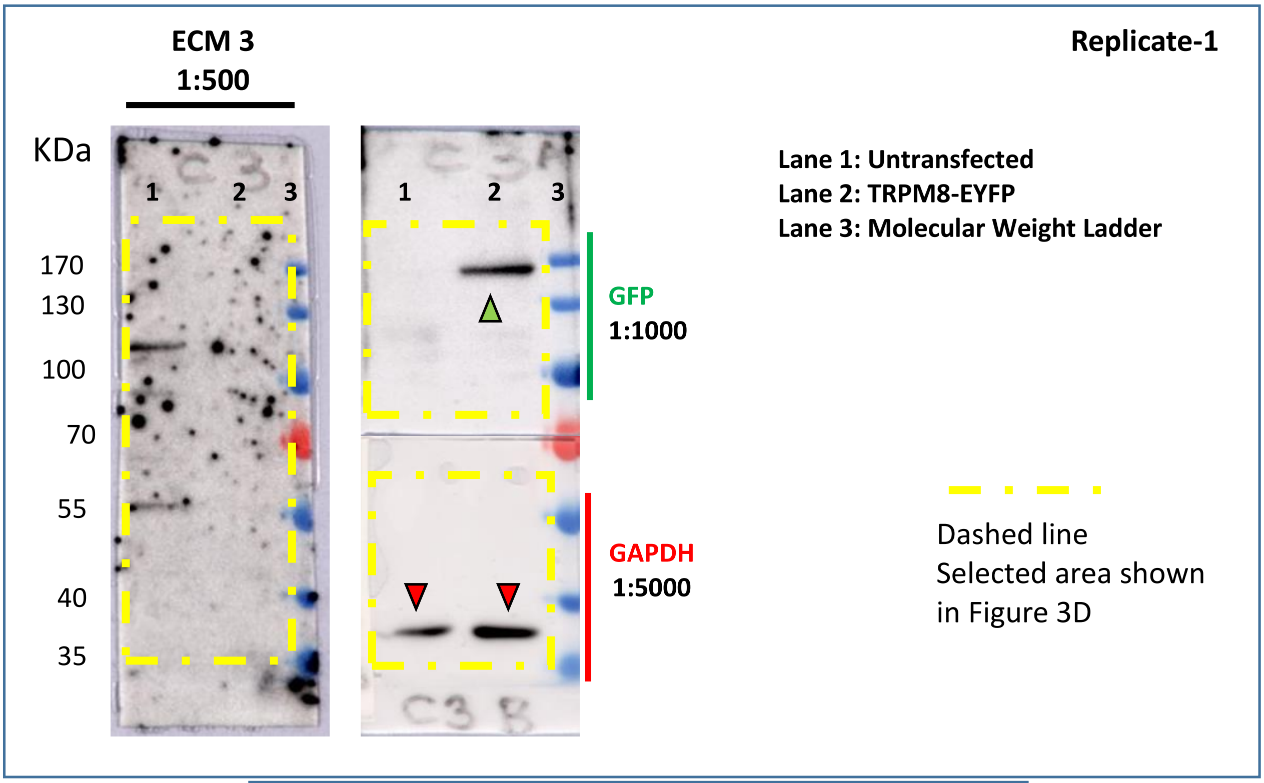
ECM 3
1:500
Replicate-1
KDa
1 2 3
170
130
100
70
55
40
35
GFP
1:1000
GAPDH
1:5000
Lane 1: Untransfected
Lane 2: TRPM8-EYFP
Lane 3: Molecular Weight Ladder
Dashed line
Selected area shown
in Figure 3D

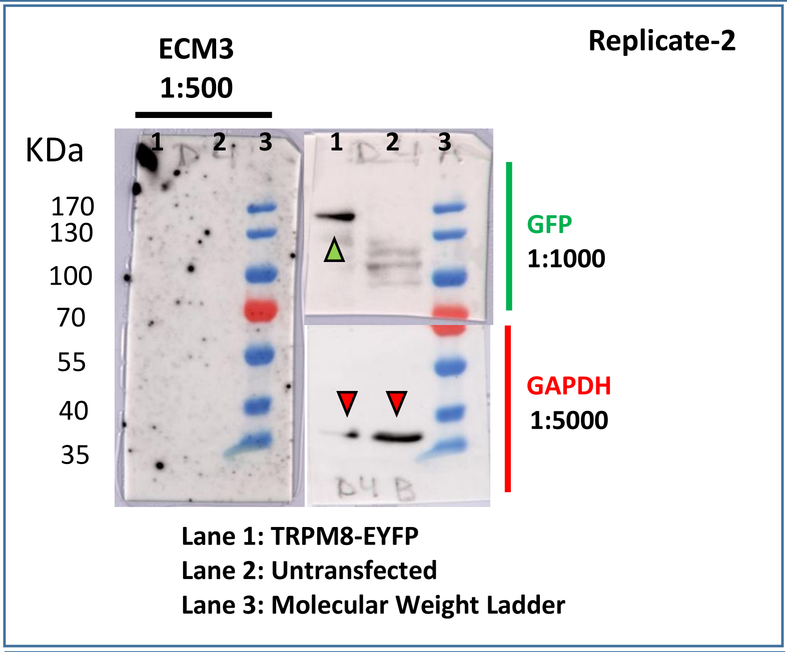
ECM3
1:500
Replicate-2
KDa
1 2 3
170
130
100
70
55
40
35
GFP
1:1000
GAPDH
1:5000
Lane 1: TRPM8-EYFP
Lane 2: Untransfected
Lane 3: Molecular Weight Ladder

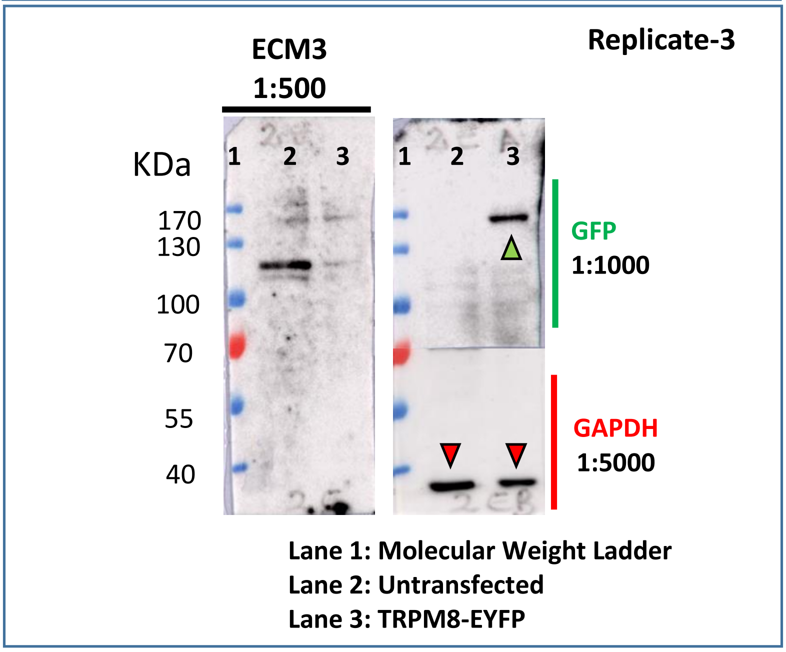
ECM3
1:500
Replicate-3
KDa
1 2 3
170
130
100
70
55
40
GFP
1:1000
GAPDH
1:5000
Lane 1: Molecular Weight Ladder
Lane 2: Untransfected
Lane 3: TRPM8-EYFP

### Origene1

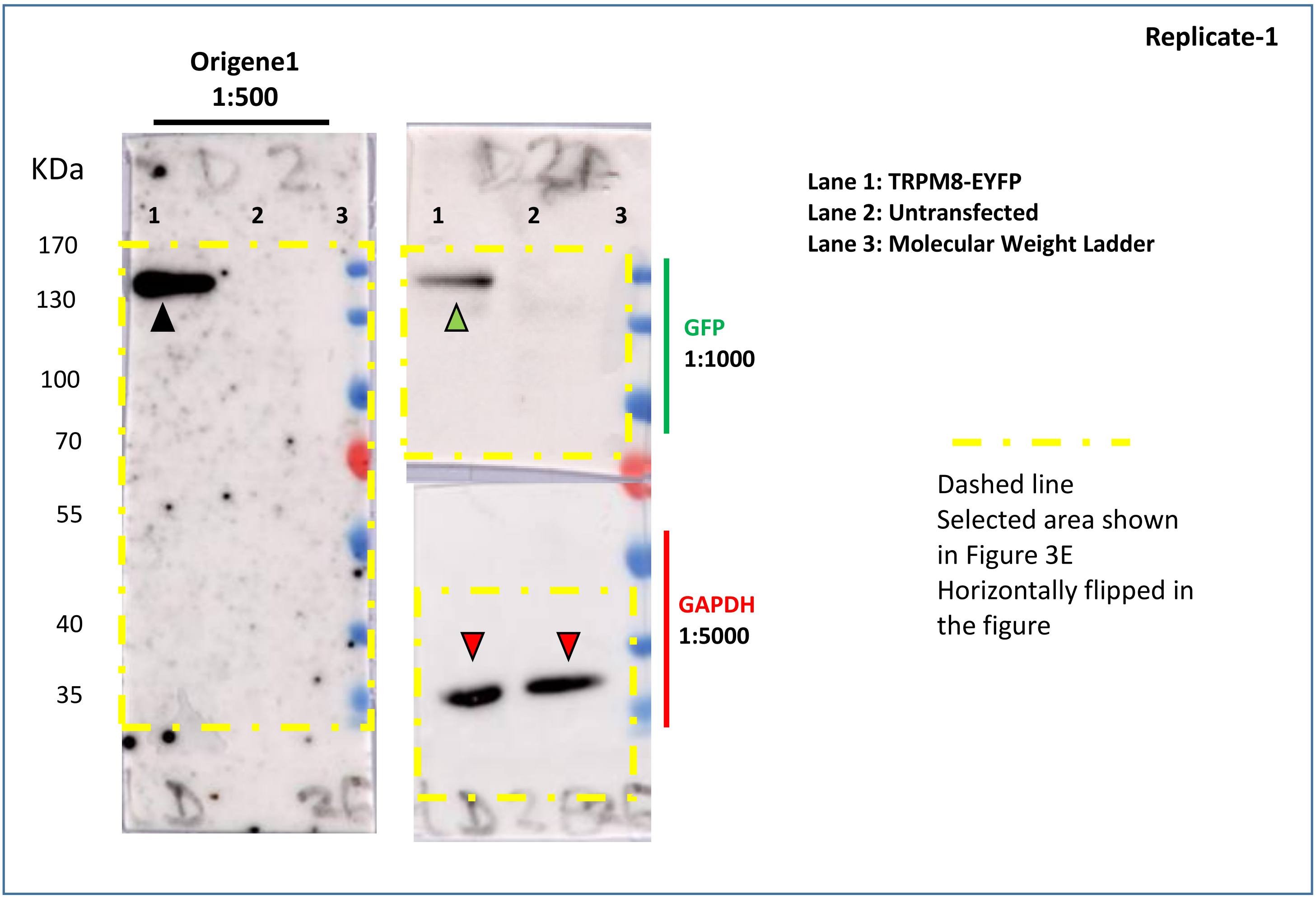

Replicate-1
Origene1
1:500
KDa
1 2 3
170
130
100
70
55
40
35
1 2 3
GFP
1:1000
GAPDH
1:5000
Lane 1: TRPM8-EYFP
Lane 2: Untransfected
Lane 3: Molecular Weight Ladder
Dashed line
Selected area shown
in Figure 3E
Horizontally flipped in
the figure

### Origene2

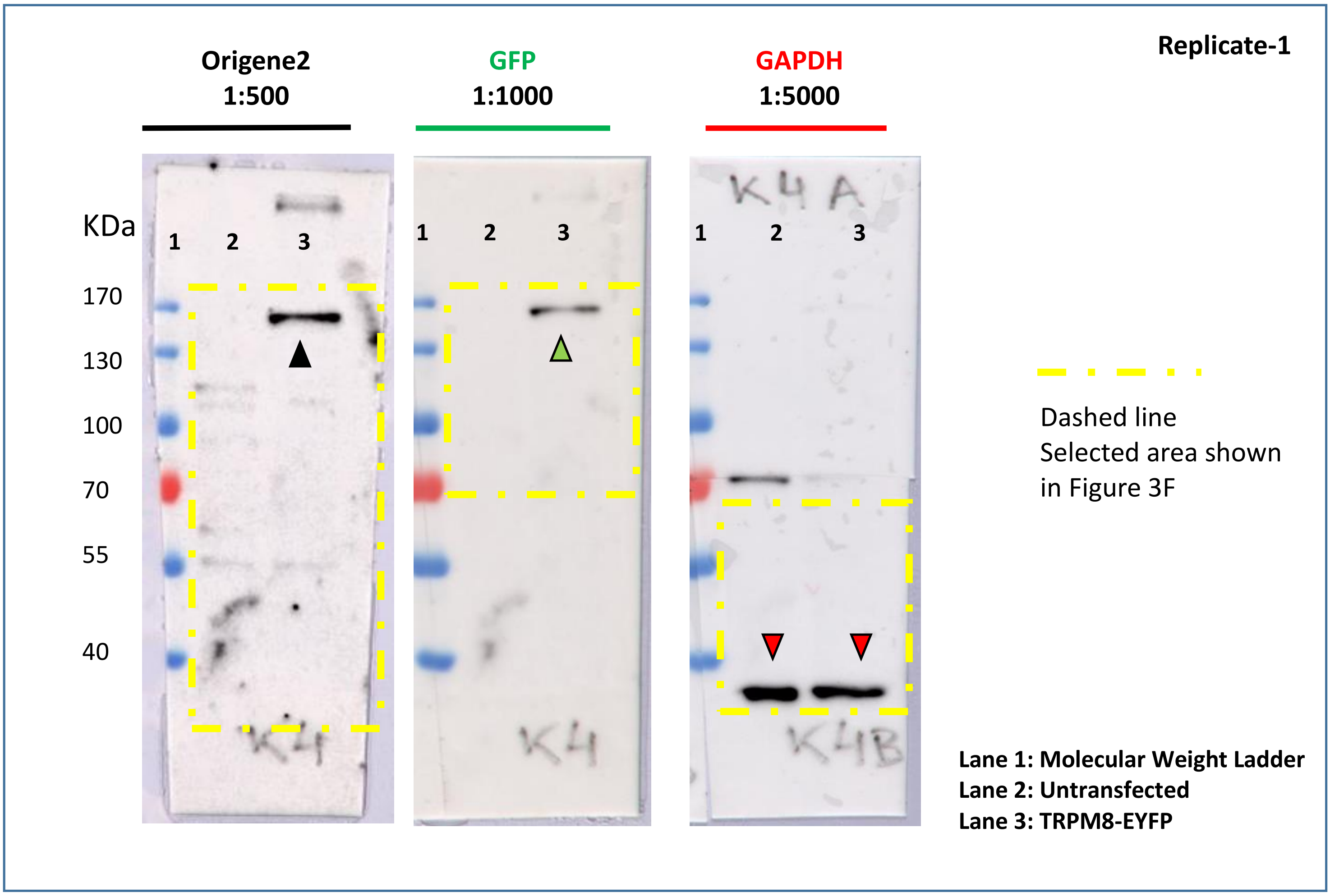


**FIGURE S4. Uncropped western blot membranes.** Full uncropped membranes are shown for each antibody. The regions shown in Figure 3 are indicated with the dashed line frames. Replicates are provided for antibodies that did not produce a TRPM8 band. Three replicates were done in all cases.
